# Harnessing Random Peptide Mixtures to Combat Multidrug-Resistant Fungal Infections

**DOI:** 10.1101/2024.09.01.610663

**Authors:** John Adeoye, Yael Belo, Marina Campos Rocha, Hilla Hayby, Aygun Israyilova, Zvi Hayouka, Neta Shlezinger

## Abstract

Invasive fungal infections are associated with high mortality and are increasingly difficult to treat due to a limited antifungal arsenal and the rapid emergence of drug resistance. Novel therapeutic strategies that combine potent antifungal activity, low host toxicity, *in vivo* stability, and a reduced propensity for resistance development are urgently needed. Antimicrobial peptides (AMPs) stand out as a promising class of compounds to combat antimicrobial resistance. Leveraging the unique properties of AMPs, we previously developed a novel approach to synthesize random peptide mixtures (RPMs) with robust bactericidal activity against drug-resistant bacteria. Here, we evaluate the antifungal potential of RPMs and demonstrate species-dependent, broad-spectrum activity of FK20 (L-phenylalanine–L-lysine, 20-mer) against major human fungal pathogens, including *Candida* spp., *Cryptococcus neoformans*, and *Aspergillus fumigatus*, with particularly high potency against the multidrug-resistant pathogen *Candida auris*. Mechanistic analyses revealed rapid membrane and cell wall disruption accompanied by intracellular penetration, consistent with membrane-active antifungal activity. Importantly, experimental evolution assays demonstrated a markedly reduced capacity for resistance development in *C. auris*. FK20 inhibited biofilm formation and displayed substantial activity against mature, pre-formed biofilms, both alone and synergistically in combination with caspofungin. Finally, FK20 showed significant therapeutic efficacy in a murine model of systemic candidiasis. Collectively, these findings establish RPMs as a versatile antifungal platform with broad-spectrum activity, biofilm efficacy, and a low resistance footprint, highlighting their promise as a novel therapeutic strategy against drug-resistant fungal infections.

**Importance:** The rising prevalence of invasive fungal infections, particularly among immunocompromised individuals, has become a critical public health concern. However, antifungal drug development has not kept pace with this growing need, and treatment options remain limited to a small number of drug classes. The emergence of multidrug-resistant fungal pathogens, such as *Candida auris*, further exacerbates this crisis by reducing the efficacy of existing therapeutics and increasing the risk of treatment failure. In this study, we evaluate the antifungal potential of FK20, a random peptide mixture (RPM) composed of L-phenylalanine and L-lysine. FK20 displays potent activity against *C. auris* and other clinically relevant human fungal pathogens, impairs biofilm formation, and exhibits synergy with caspofungin. Importantly, FK20 limits the emergence of resistance and demonstrates therapeutic efficacy in a murine model of systemic candidiasis. These findings establish RPMs as a promising new class of antifungals with broad-spectrum activity and clinical potential against drug-resistant fungal infections.

## Introduction

The surge in antimicrobial resistance poses a significant threat to public health (1–4). While attention has predominantly centered on pan-resistant bacteria, mounting apprehension surrounds multidrug-resistant fungal pathogens, notably *Candida auris* (5). This emerging pathogen poses a significant global health threat due to its rapid worldwide spread and intrinsic multidrug resistance (6,7). *C. auris* has been recognized as a serious global health threat by both national and international public health agencies (8,9), underscoring the necessity for additional therapeutic options to address drug-resistant fungal infections.

*C. auris* was initially identified in 2009 in a Japanese hospital, where it was isolated from the external ear canal of a patient (10). Since then, genomic analysis has shed light on the concurrent emergence of distinct lineages, categorizing them into six major geographical clades across six continents over the past ∼400 years (11–14). Its rapid transmission within clinical settings is attributed to its remarkable long-term persistence on surfaces, including human skin, as well as the widespread prevalence of drug-resistant isolates (15). Hospital-acquired *C. auris* infections, most commonly presenting as candidemia, occur predominantly in patients previously colonized with *C. auris*, most frequently on the skin, which serves as a persistent reservoir and a probable portal facilitating transmission and invasive disease, with reported mortality rates ranging from 30% to 60% (15,16). Risk factors that increase susceptibility to *C. auris* infections are shared with other fungal infections and include iatrogenic, primary, or acquired immunodeficiency, prolonged antibiotic regimens, prolonged hospitalization, and invasive medical procedures (17).

*C. auris* is classified as a ‘superbug fungus’, posing an increasingly concerning threat to human health due to varying levels of intrinsic resistance. Approximately 5-10% of isolates exhibit resistance to all three major antifungal drug classes commonly used in clinical settings (18,19). It is evident that over 80% of *C. auris* isolates are resistant to at least one azole antifungal, most commonly fluconazole (20), and many strains exhibit notably elevated minimum inhibitory concentrations (MIC) for polyenes such as amphotericin B (21–23). For these reasons, echinocandins such as caspofungin, are used as front-line therapeutics against systemic candidemia caused by *C. auris* (24–26). However, isolates with reduced susceptibility to one or more echinocandins have also been reported (14, 25, 26) (20,27,28). Similar to other *Candida* species, *C. auris* employs diverse drug resistance mechanisms, including point mutations or chromosomal rearrangements affecting drug targets, as well as upregulation of efflux pumps. Collectively, the co-occurrence of multidrug resistance, rapid global emergence, and high mortality rates designates *C. auris* as a pathogen of significant clinical concern and complexity, highlighting the urgent need for novel antifungal strategies.

Current antifungal therapies are further limited by a narrow spectrum of drug classes, toxicity concerns, drug–drug interactions, and the emergence of resistance during prolonged treatment. In this context, alternative therapeutic paradigms that exploit fundamentally different modes of action are increasingly sought. One such strategy involves antimicrobial peptides (AMPs), an evolutionarily conserved component of innate immunity, including naturally occurring peptides such as defensins, cathelicidins, and histatins that are integral to innate immune systems across diverse organisms and exhibit broad-spectrum activity against bacteria, fungi, viruses, and parasites (29–32). AMPs are typically short amphipathic peptides rich in cationic and hydrophobic residues, that exert antimicrobial activity primarily through interactions with microbial membranes, leading to membrane destabilization and cell death (33–35). Several endogenous and synthetic AMPs, including human defensins and antifungal proteins (AFPs) from filamentous fungi, have demonstrated antifungal activity against *Candida*, *Aspergillus*, and *Cryptococcus* species, underscoring the therapeutic promise of peptide-based antifungals (36–38).

Building on the fundamental principles underlying AMP activity, we previously developed a novel class of antimicrobial agents termed Random Peptide Mixtures (RPMs). RPMs were originally conceived as a synthetic alternative to natural antimicrobial peptides, aiming to overcome key limitations that have hindered AMP clinical translation, including high production costs, and the rapid evolution of resistance associated with uniform, well-defined peptide sequences (39–41). RPMs are generated by solid-phase synthesis in which, at each coupling step, a defined mixture of one cationic and one hydrophobic amino acid is incorporated, resulting in a highly diverse ensemble of peptides with identical chain length, stereochemistry, and overall composition (42). This combinatorial design produces molecular heterogeneity that preserves antimicrobial potency while reducing selective pressure for resistance. RPMs have been shown to exhibit robust antimicrobial efficacy against both Gram-positive and Gram-negative bacteria *in vitro* and in multiple mouse infection models, including methicillin-resistant *Staphylococcus aureus* and *Acinetobacter baumannii* (43–46). Importantly, prior studies have demonstrated favorable safety profiles and limited host toxicity *in vivo*, supporting the translational potential of this approach.

Here, we present for the first time the antifungal activity of RPMs against the multidrug resistant fungal pathogen *C. auris*. We demonstrate potent *in vitro* and *in vivo* efficacy of the RPMs, assess resistance development and collateral sensitivity, and explore synergistic interactions with established antifungals, thereby positioning RPMs as promising candidates for future antifungal development.

## Results

### Random Peptide Mixtures Demonstrate Strong Antifungal Activity against *C. auris In Vitro*

To elucidate the antifungal efficacy of random peptide mixtures (RPMs) against *C. auris*, killing assays were conducted using the clinical isolate B11117 to compare RPMs based on their amino acid composition, chain length and stereochromy. Among the four tested RPMs composed of cationic and hydrophobic amino acids—FK20 (L-Phenylalanine–L-Lysine, 20-mer), LK20 (L-Leucine–L-Lysine, 20-mer), IK20 (L-Isoleucine–L-Lysine, 20-mer), and IK10 (L-Isoleucine-L-Lysine, 10-mer) —FK20 exhibited superior antifungal activity against *C. auris* (Figure 1A). Given its minimal antifungal activity against *C. auris*, IK20 was selected as the negative control RPM. To further explore RPM optimization, the antifungal efficacy of FK RPMs with varying chain lengths was assessed (Figure 1B). The 5 and 10-mer FK RPMs did not exhibit any discernible antifungal properties, whereas the 30-mer FK RPM showed marginal activity. In contrast, FK20 RPM demonstrated the highest antifungal activity, consistent with its previously reported broad antimicrobial potency and membrane-active properties (47). Next, the impact of FK RPM stereochemistry was assessed (Figure 1C). Three stereoisomers were compared: FK20 (L-Phenylalanine−L-Lysine, 20-mer), FdK20 (L-Phenylalanine−D-Lysine, 20-mer), and dFdK20 (D-Phenylalanine−D-Lysine, 20-mer). The homochiral RPMs, FK20 and dFdK20, exhibited robust antifungal activity against *C. auris*, whereas the heterochiral RPM FdK20 displayed comparatively lower activity levels. Based on these findings, homochiral FK20 RPM was identified as the most effective candidate for combating *C. auris* and was selected for all subsequent experiments.

**Figure 1:**
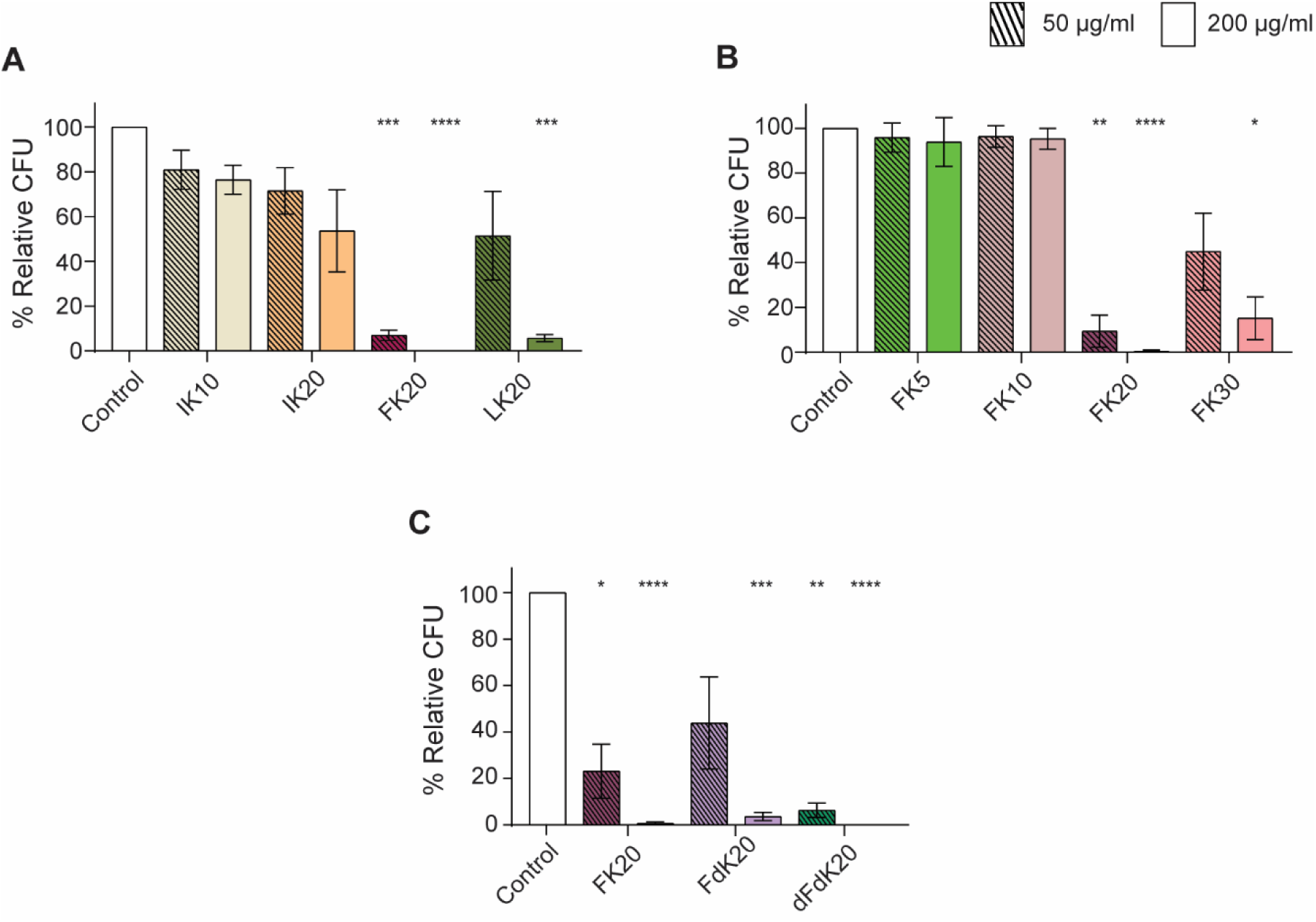
FK20 RPM exhibits potent antifungal activity against *C. auris*. *C. auris* cells (1 x 10^6^ cells/ml) were incubated with either 50 μg/ml (striped bars) or 200 μg/ml (solid bars) of the indicated RPMs for 45 min at 37°C with shaking (200 rpm) in PBS. Fungicidal activity was assessed based on: (**A)** amino acid composition (IK10, IK20, FK20 and LK20), (**B)** peptide chain length (5, 10, 20, and 30-mer FK), (**C)** Stereochemistry (FK20, FdK20, and dFdK20). Fungal viability was quantified by CFU enumeration and expressed as a percentage of untreated control. Data represents the mean ± SEM of three biologically independent experiments. Statistical significance was determined using students t-test, comparing each treatment to the untreated control (set to 100% for each biological replicate). Significance levels are indicated as *p <0.05, **p < 0.01, ***p<0.001 and ****p < 0.0001; values without symbols are not significantly different from the control.

### FK20 RPM displays species-dependent antifungal activity across major fungal pathogens

To determine whether FK20 RPM activity extends beyond *Candida* spp., we evaluated its antifungal efficacy against additional clinically relevant fungal pathogens representing distinct phylogenetic and structural classes: *Cryptococcus neoformans* and *Aspergillus fumigatus*. FK20 RPM activity was compared to IK20 RPM and to the reference antifungals caspofungin and amphotericin B (Figure 2A, Figure S1 A-B).

**Figure 2:**
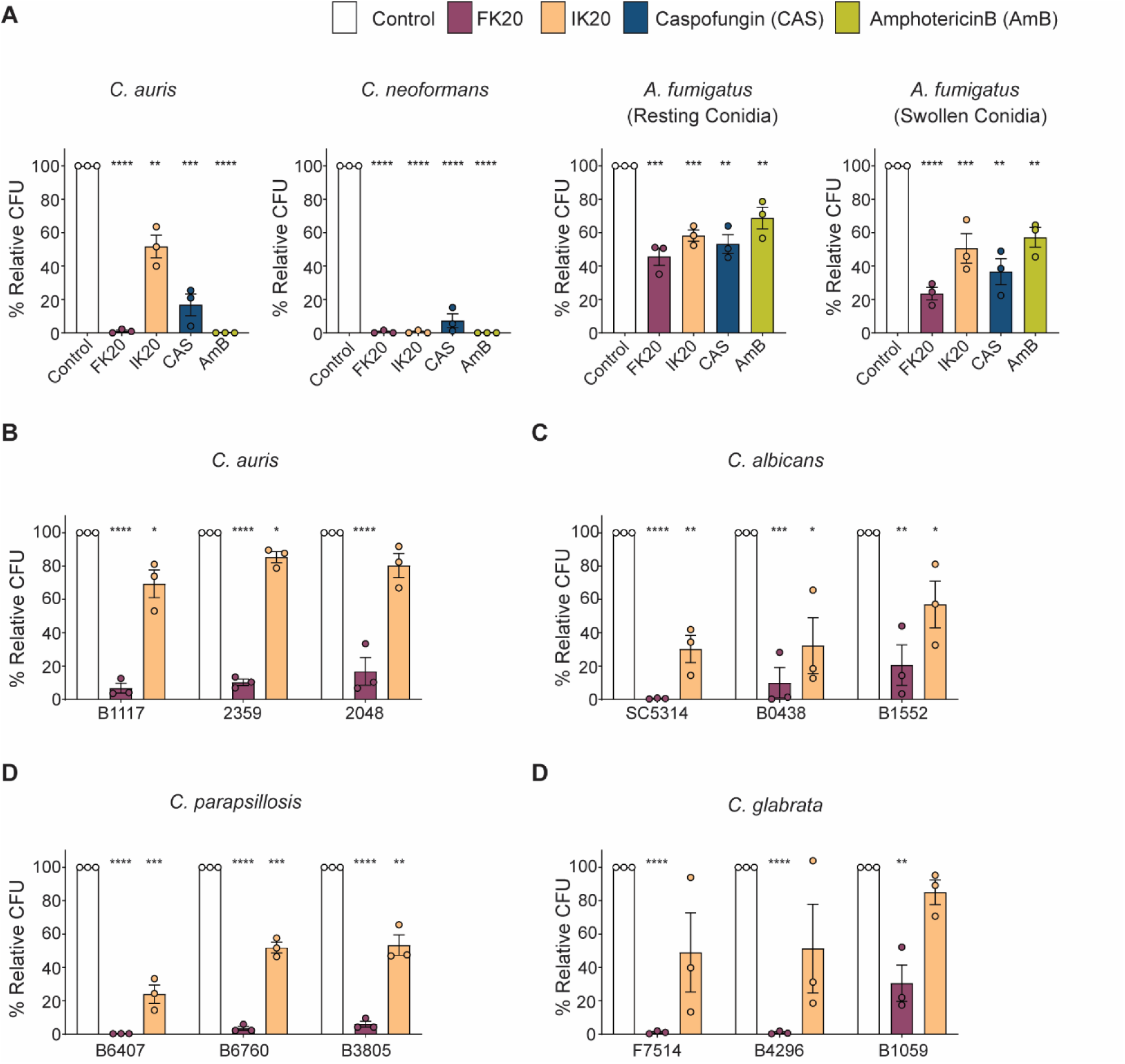
FK20 potency varies across major fungal pathogens of clinical relevance. (**A)** *C. auris* and *C. neoformans* yeast cells, as well as *A. fumigatus* conidia (resting and swollen), were incubated with 200µg/ml FK20 or IK20 RPMs, 2µg/ml Caspofungin, 2µg/ml Amphotericin B, or PBS as a control for 5h at 37°C, followed by CFU determination. **(B-E)** Cells of *C. auris* **(B),** *C. albicans* **(C),** C. *parapsillosis* **(D),** and *C. glabrata* **(E)** were treated with 200µg/ml FK20 or IK20 RPMs, or PBS as a control, for 45 min at 37°C, followed by CFU determination. Clinical isolates included *C. auris* skin-derived isolates (1117, 2359, 2048); *C. albicans* blood-derived reference strain SC5314 and blood-derived isolates (B0438, B1552); *C. parapsillosis* blood-derived isolates (B6407, B6760, B3805); and *C. glabrata* isolates derived from peritoneal fluid (F7514) and blood (B4296, B1059). Fungal viability is presented as CFUs expressed as a percentage of untreated control. Data represents the mean ± SEM of three biologically independent experiments. Statistical significance was determined using student’s t-test, comparing each treatment to the untreated control (set to 100% for each biological replicate). Significance levels are indicated as *p <0.05, **p < 0.01, ***p<0.001 and ****p < 0.0001; values without symbols are not significantly different from the control.

FK20 RPM exhibited potent fungicidal activity against *C. neoformans*, comparable to its activity against *C. auris*. In contrast, FK20 retained significant but reduced activity against *A. fumigatus*, achieving 77% killing of swollen conidia and 54% killing of resting conidia. IK20 RPM showed minimal fungicidal activity against *C. auris* and *A. fumigatus* (48% killing against *C. auris*, and 42% and 50% killing against resting and swollen *A. fumigatus* conidia, respectively), but retained high activity against *C. neoformans* (99% killing).

In comparison, standard antifungal agents displayed potent activity against the yeast pathogens *C. auris* and *C. neoformans*, but markedly reduced efficacy against the filamentous fungus *A. fumigatus* under the tested conditions. Amphotericin B exhibited strong activity against *C. auris* and *C. neoformans* (100% killing for both), but substantially weaker activity against *A. fumigatus* (31% and 43% killing against resting and swollen conidia, respectively). Caspofungin showed limited efficacy against *A. fumigatus* (47% and 63% killing against resting and swollen conidia, respectively) and moderate activity against *C. neoformans* and *C. auris* (93% and 83% killing, respectively), both lower than that observed for FK20.

Together, these data reveal marked species-specific differences in susceptibility to FK20 RPM, with highest potency against *C. auris* and *C. neoformans* and reduced—yet clearly detectable—activity against *A. fumigatus*.

To further delineate FK20 RPM activity within the *Candida* genus, its antifungal efficacy was evaluated against clinical isolates of *C. auris* and other clinically relevant *Candida* species, including *C. albicans*, *C. parapsilosis*, and *C. glabrata*. Cells were treated with 200µg/ml FK20, IK20 RPM (control), or PBS. FK20 RPM demonstrated strong antifungal activity across all tested *Candida* species, with near-complete killing of all *C. parapsilosis* isolates and the majority of *C. auris*, *C. albicans*, and *C. glabrata* isolates (Figure 2B–E). While the magnitude of killing varied between species and individual isolates, FK20 RPM consistently achieved substantial reductions in fungal viability across all tested clinical isolates. Together with its activity against non-*Candida* fungal pathogens, these findings establish FK20 RPM as a broad-spectrum antifungal agent whose efficacy is modulated by species– and isolate-specific properties.

### FK20 RPM Disrupts *Candida auris via* Cell Wall Penetration, Membrane Damage, ROS Induction, and Membrane Depolarization

To elucidate the mechanisms underlying FK20 RPM’s antifungal activity, its interaction with *C. auris* cells was examined using a combination of fluorescence microscopy, flow cytometry, and membrane integrity assays. Because RPMs are designed to mimic the physicochemical properties of antimicrobial peptides rather than engage a specific molecular target, these experiments were aimed at defining the cellular consequences of FK20 exposure rather than identifying a single receptor or uptake pathway. The ability of FK20 RPM to associate with and penetrate *C. auris* cells was assessed using N-terminally 5(6)-carboxyfluorescein-labeled FK20 RPM. *C. auris* cells incubated with fluorescently labeled FK20 for 45 min and stained with Calcofluor White were visualized by confocal microscopy. Fluorescein-labeled FK20 (green) was detected within the boundaries of the Calcofluor White stained cell wall (blue), as confirmed by Z-stack imaging and three-dimensional reconstruction (Figure 3A; Supplementary Movie 1). Importantly, fluorescein labeling did not impair FK20 antifungal activity, as labeled FK20 retained fungicidal potency comparable to unlabeled FK20, as assessed by CFU enumeration (Fig. 3B). Consistent with this retained activity, fluorescein-labeled FK20 induced slightly increased membrane permeabilization relative to unlabeled FK20, as reflected by enhanced propidium iodide uptake, a phenomenon previously reported for N-terminally modified antimicrobial peptides and attributable to augmented membrane interactions rather than a distinct mechanism of action (Fig. 3C) (48). To exclude nonspecific dye uptake or fluorescence artifacts, a series of specificity controls were included. Free fluorescein did not accumulate in *C. auris* cells under any condition tested, either when applied alone or when co-incubated with unlabeled FK20, indicating that intracellular fluorescence required peptide conjugation and was not driven by membrane damage–mediated dye entry (Fig. 3A). In contrast, fluorescein-labeled FK20 showed robust cellular association, consistent with its retained antifungal activity. Fluorescein-labeled IK10 and LK20 RPMs—peptides displaying only minimal antifungal activity—exhibited markedly reduced cellular association under identical labeling and imaging conditions (Fig. 3A). Flow cytometric quantification confirmed significantly higher uptake of FK20 relative to IK10 and LK20, demonstrating that intracellular accumulation correlates with antifungal potency rather than nonspecific membrane disruption or dye uptake (Fig. 3D).

**Figure 3:**
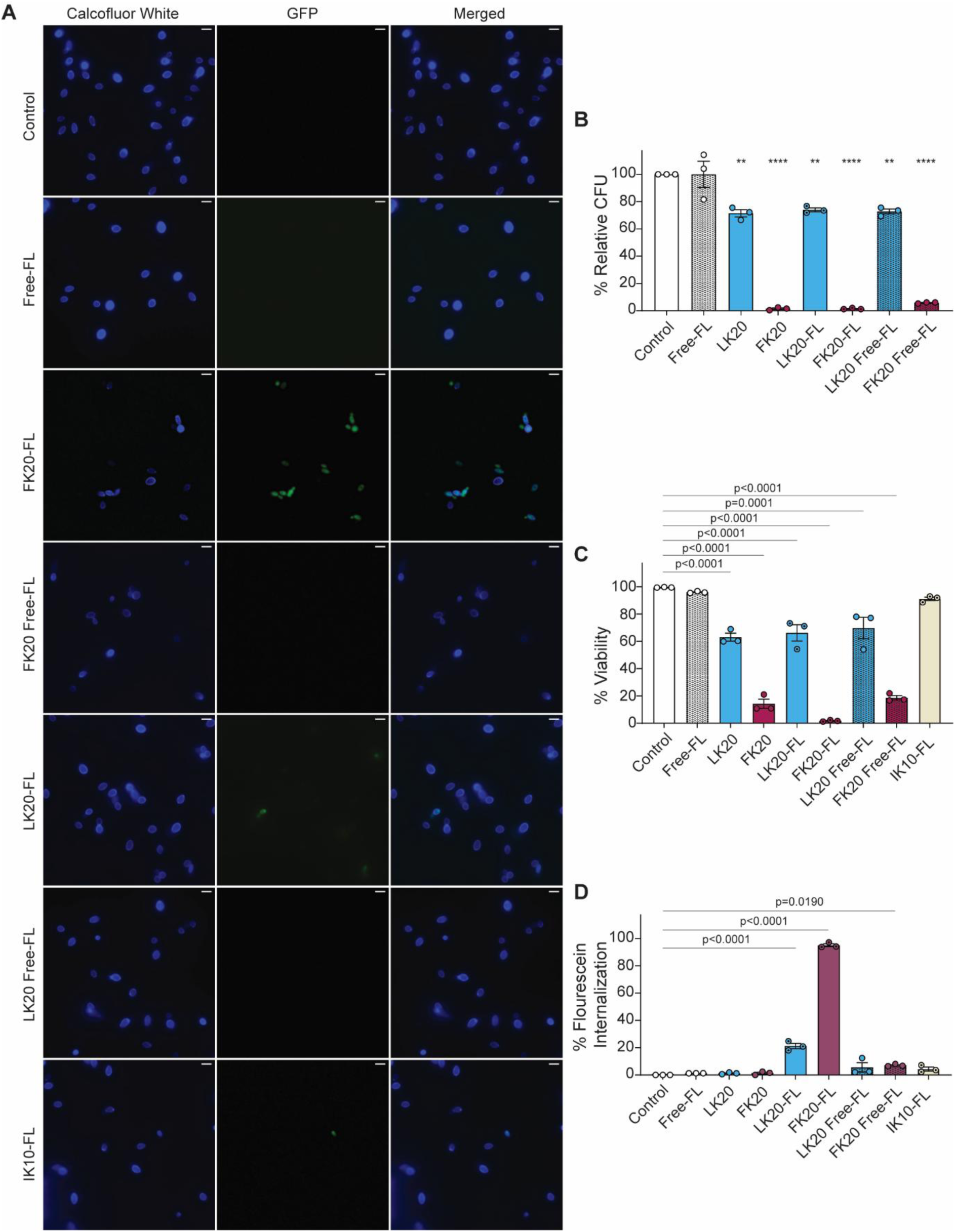
FK20 RPM penetrates *C. auris* cells. (**A)** *C. auris* cells (5 x 10^7^ cells/ml) were incubated with 200ug/ml fluorescein-conjugated FK20, LK20 and 1K10 RPMs for 45 mins at 37°C in PBS. Controls included untreated cells, cells treated with free fluorescein alone, and cells treated with free fluorescein co-incubated with unlabeled FK20 or LK20. Cells were stained with Calcofluor White and visualized by fluorescence microscopy (100x, scale bar = 5 um). Representative images from three independent experiments are shown. (**B-C)** Antifungal activity of fluorescein labeled RPMs determined by **(B)** CFU enumeration and **(C)** PI staining. *C. auris* cells (1 x 10^6^ cells/ml) were treated as above; data represents mean CFU or percentage PI-negative (viable) cells from three independent experiments performed in triplicates. (**D)** Internalization of fluorescein-conjugated RPMs assessed by flow cytometry. Data represents mean percentage of FITC-positive cells from three independent experiments performed in triplicates. Statistical significance was determined using student’s t-test, comparing each treatment to the untreated control (set to 100% for each biological replicate). Significance levels are indicated as *p<0.05, **p<0.01, ***p<0.001 and ****p<0.0001; values without symbols are not significantly different from the control.

To investigate FK20-induced surface morphological changes, scanning electron microscopy (SEM) was performed on *C. auris* cells treated with 200 µg/ml of FK20 RPM stereoisomers (Fig. 4A). Homochiral RPMs (FK20 and dFdK20) induced pronounced surface disruption and extensive cellular aggregation compared to heterochiral FdK20 RPM and untreated controls. These findings indicate that homochirality enhances physicochemical interactions with the fungal surface, leading to more effective membrane destabilization. Membrane integrity was further assessed by measuring propidium iodide (PI) uptake using flow cytometry. FK20 treatment resulted in a concentration-dependent increase in PI-positive cells, confirming progressive membrane permeabilization (Figure 4B). Similar results were observed using Ghost Dye, an amine-reactive viability dye (Figure S2). Because RPM-induced membrane damage enables dye entry, intracellular fluorescence is expected to follow loss of membrane integrity rather than precede it, consistent with a membrane-disruptive mechanism of action.

**Figure 4:**
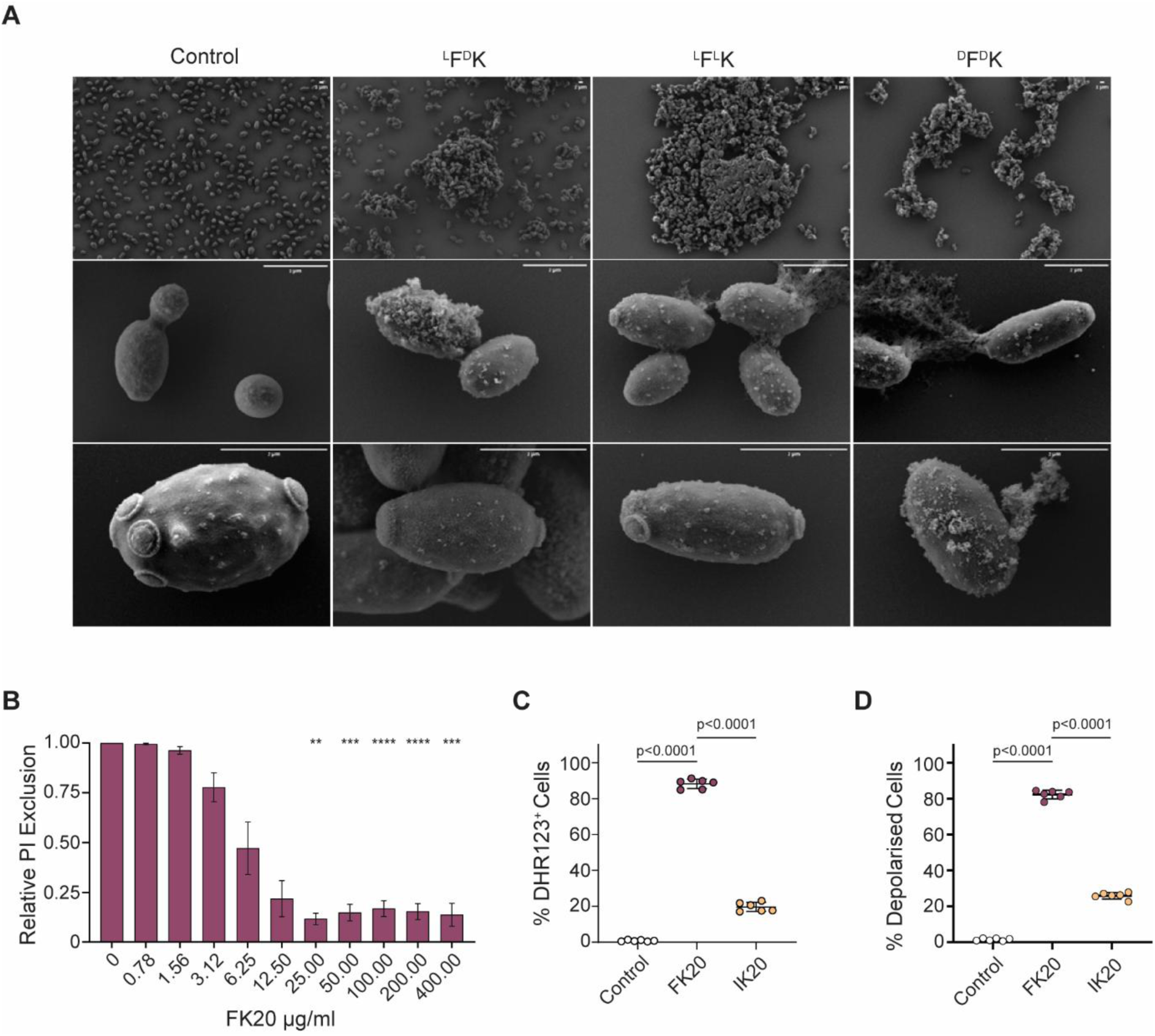
*C. auris* cell surface after interaction with 20-mer FK RPMs. (**A)** Scanning electron microscopy of untreated *C. auris* and cells treated with 200 μg/ml of 20-mer FK20, FdK, or dFdK using JEOL JSM-7800f scanning electron microscope at 20 kV (scale bar = 1μm). Representative micrographs from three independent experiments are shown. (**B-D)** FK20 induces membrane damage, ROS generation, and membrane depolarization in *C. auris* cells. Cells (1 x 10^6^ cells/ml) were treated with FK20 for 45 min at 37°C in PBS. Cells were analyzed using flow cytometry. **(B)** Dose-dependent membrane damage induced was determined by incubating cells with FK20 at concentrations ranging from 0-400ug/ml. Membrane integrity was assessed by PI uptake into the cells (flow cytometry). Data are presented as PI excluded cells (PI negative) relative to untreated control. (**C)** Intracellular Reactive Oxygen Species (ROS) generation was assessed by staining *C. auris* cells with DHR 123 after treatment with FK20. **D.** Membrane depolarization was assessed using DIBAC_4_(5) flowing FK20 treatment. Data represent mean ±SEM from three independent experiments performed in triplicates. Statistical significance determined by one-way ANOVA with Turkey’s multiple comparisons test. Significance levels are indicated as *p<0.05, **p<0.01, ***p<0.001 and ****p<0.0001; values without symbols are not significantly different from the control.

Given that membrane-active antimicrobials often induce secondary cellular stress responses (49,50), intracellular reactive oxygen species (ROS) ROS production was measured using dihydrorhodamine 123 (DHR123) (Figure 4C). FK20-treated *C. auris* exhibited significantly increased ROS levels relative to controls. In parallel, membrane depolarization was assessed using Bis-(1,3-dibutylbarbituric acid) pentamethine oxonol (DiBAC₄(5)), revealing a marked increase in fluorescence following FK20 exposure, indicative of collapse of membrane potential (Figure 4D). Collectively, these data demonstrate that FK20 RPM exerts antifungal activity through a physicochemical mechanism dominated by membrane disruption, loss of membrane potential, ionic imbalance, and downstream oxidative stress, rather than through engagement of a specific intracellular target.

### FK20 RPM Does Not Induce Resistance in *C. auris* and Exhibits Collateral Sensitivity in Caspofungin-Resistant Strains

The rapid evolution of drug resistance in pathogens is a primary cause of treatment failure, often diminishing the efficacy of new antifungals shortly after market entry. Evaluating a drug candidate’s resistance potential is therefore critical. Our previous studies on the antibacterial activity of RPMs demonstrated that bacteria were unable to develop resistance, suggesting a promising low-resistance potential for this class of compounds (51,52). To investigate the potential of *C. auris* to acquire resistance against FK20 RPM, *in vitro* experimental evolution was conducted following established protocols with minor modifications (53) (Figure 5A). The FK20 RPM MIC_50_, defined as the drug concentration inhibiting growth by at least 50% relative to drug-free control, was first determined (Figure S3A), resulting in values of 23.80±7.09 µg/ml at 24h, 54.16±11.67 µg/ml at 48h, and 104.10±19.82 µg/ml at 72h. Three parallel *C. auris* lineages were evolved in RPMI-1640 media containing FK20 RPM (100 µg/ml, MIC₅₀), caspofungin (2.0 µg/ml, MIC₅₀), or no drug (ND-Evol) through ten serial passages, diluted 1:1,000 every 72 hours, resulting in the emergence of three independently evolved strains for each condition. Evolved strains were then challenged with their respective antifungal agents, and growth kinetics were measured as optical density (OD) after 72 hours to determine new MIC₅₀ values, compared to parental and ND-Evol strains. FK20-evolved strains showed no significant change in the FK20 MIC_50_ relative to parental or ND-Evol strains (Figure 5, Figure S2 B-C, Table 1). In contrast, caspofungin-evolved strains exhibited a marked increase in caspofungin MIC₅₀ to >10 µg/ml (Figure 5C, Figure S2D–E). These findings demonstrate that while *C. aur*is rapidly develops resistance to caspofungin within ten passages, it fails to evolve resistance to FK20 RPM over the same timeframe, highlighting FK20’s potential to limit resistance development.

**Figure 5.**
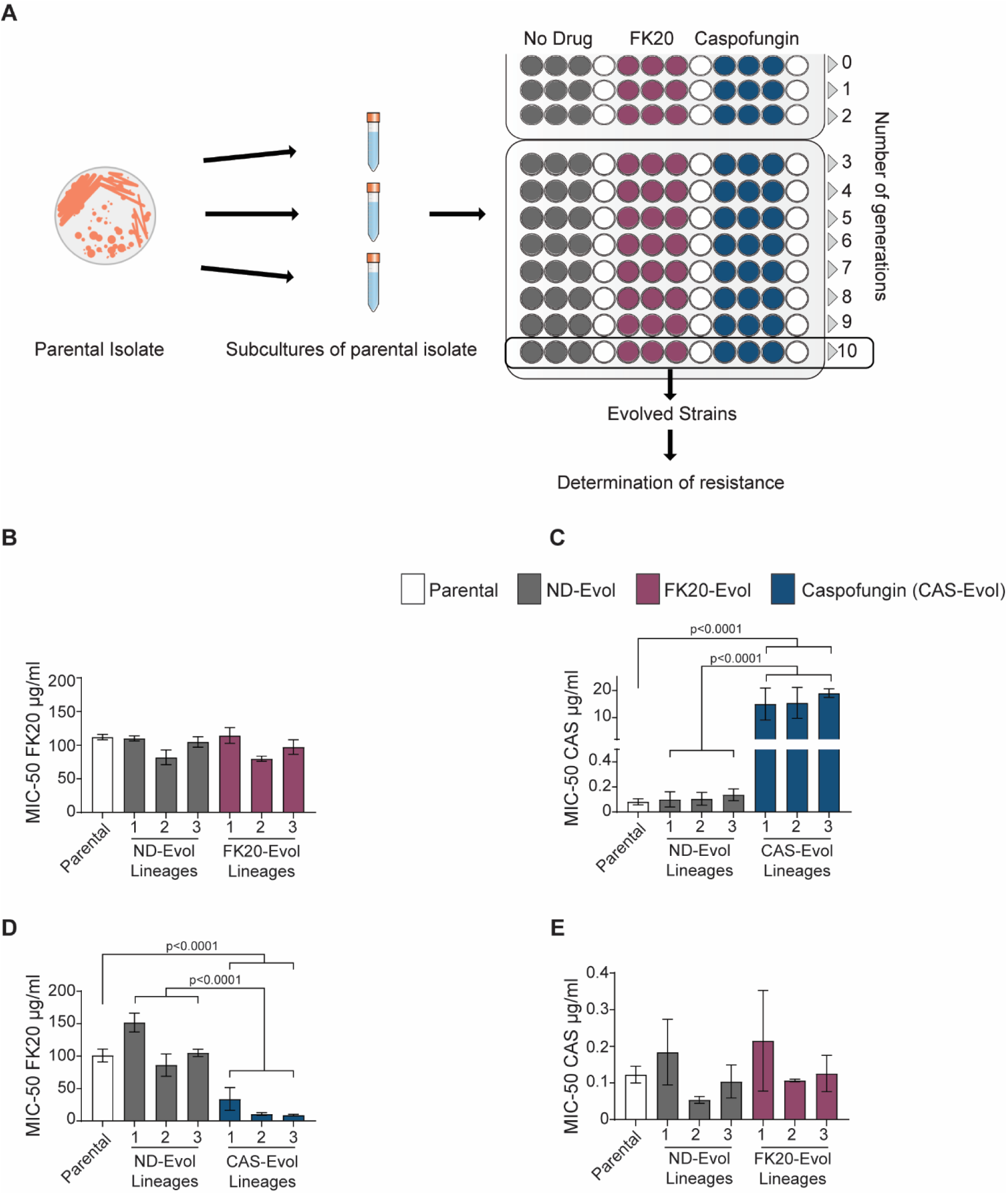
*In vitro* experimental evolution of *Candida auris*. (A) Schematic representation of the experimental evolution strategy. Parental *C. auris* cells were propagated under three conditions: without treatment, or in the presence of either FK20 or caspofungin to generate three independent evolved lineages after ten sequential passages. Following evolution, growth profiles and resistance levels were determined by calculating MIC-50 values for the parental, non-drug-evolved, and drug-evolved strains. Broth microdilution assays were performed to quantify growth by measuring optical density at OD-590 nm across a range of increasing FK20 and caspofungin concentrations and determine MIC-50. Data is presented as MIC-50 after 72 hours of incubation under the respective drug conditions. (B) MIC-50 values for FK20 in parental, non-drug-evolved, and FK20-evolved lineages. (C) MIC-50 values for caspofungin in parental, non-drug-evolved, and caspofungin-evolved lineages. (D–E) Collateral sensitivity analysis of evolved strains. (D) FK20 MIC-50 values for parental, non-drug-evolved, and caspofungin-evolved lineages. (E) Caspofungin MIC-50 values for parental, non-drug-evolved, and FK20-evolved lineages. Each data point represents the mean ± S.E. derived from three independent experiments performed in triplicate. Statistical significance determined by one-way ANOVA with Turkey’s multiple comparisons test. Values without symbols are not significantly different from the control.

**Table 1.**
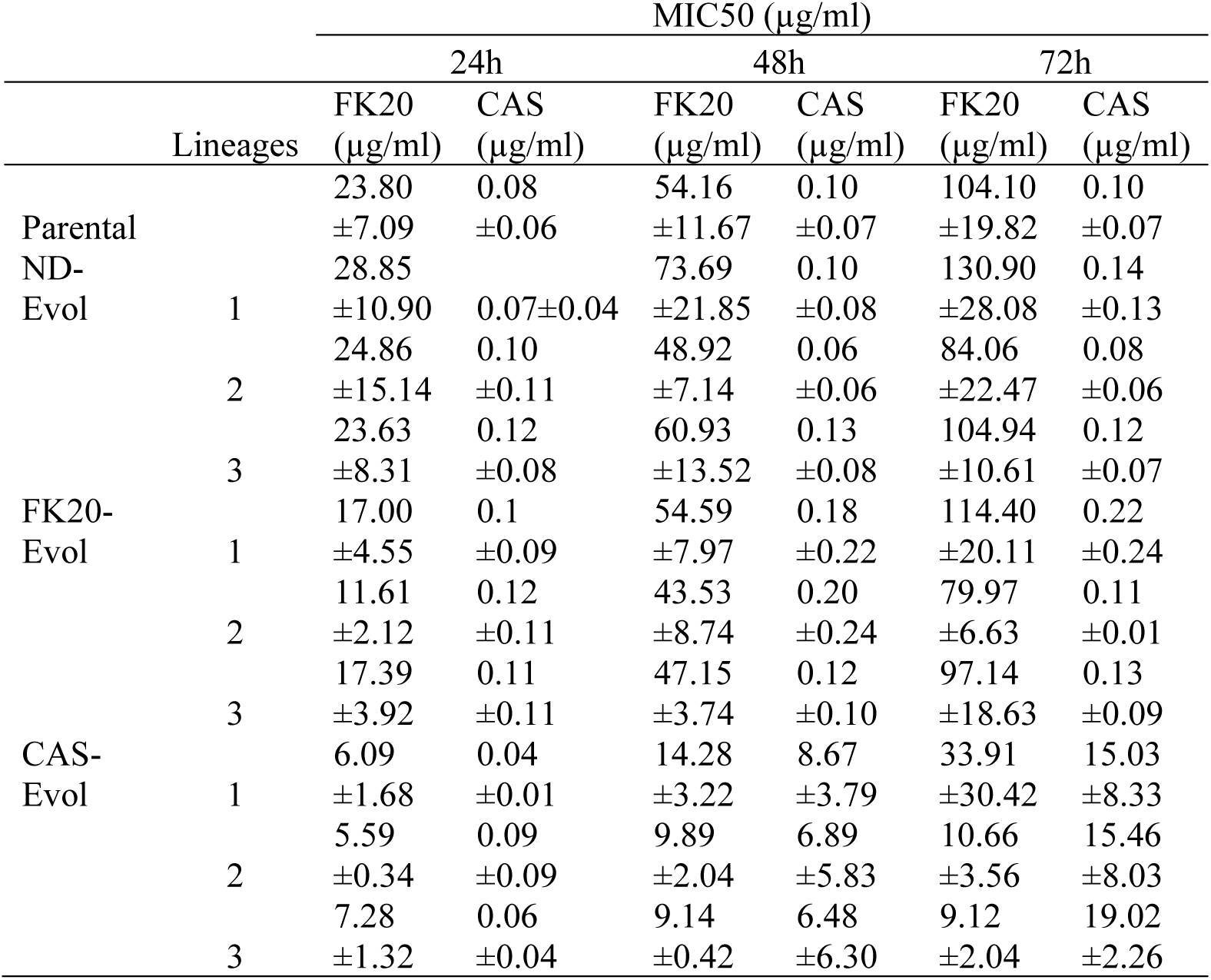
MIC_50_ profiles of parental and evolved strains over time. Minimum inhibitory concentrations (MIC_50_ ± SEM) for FK20 and caspofungin were determined for the parental strain and experimentally evolved strains (ND-Evol, FK20-Evol, and CAS-Evol; three independent lineages per evolved condition) after 24, 48, and 72 h of incubation. Values represent the mean ± SEM from three independent experiments.

Collateral sensitivity, an evolutionary trade-off where resistance to one antifungal agent increases susceptibility to another (54), was by assessing the resistance profiles of FK20-evolved and caspofungin-evolved *C. auris* strains when challenged with the reciprocal antifungal (caspofungin or FK20 RPM, respectively). FK20-evolved strains show no significant change in caspofungin MIC_50_ compared to parental or ND-Evol strains (Figure 5E, Figure S2 H-I). In contrast, caspofungin-evolved strains exhibited a marked reduction in FK20 RPM MIC-50 (Figure 5D, Figure S2 F-G), indicating collateral sensitivity to FK20 RPM. These findings suggest that while FK20 exposure does not alter caspofungin susceptibility, caspofungin resistance enhances susceptibility to FK20 RPM, a property with potential therapeutic applications.

### FK20 RPM and Caspofungin are Synergistic and Inhibit *C. auris* Biofilm Formation

Combination therapies leveraging collateral sensitivity can reduce resistance evolution, as mutations conferring resistance to one antifungal may increase susceptibility to another (55). Given the observed collateral sensitivity to FK20 RPM in caspofungin-resistant *C. auris* strains, the potential synergistic interaction between FK20 RPM and caspofungin was investigated. FK20 disrupts *C. auris* cell walls and membranes, while caspofungin inhibits the enzyme (1→3)-β-D-glucan synthase, compromising cell wall integrity (56). It was hypothesized that their combined treatment would induce synergistic cell wall damage and osmotic stress. A checkerboard assay was conducted, challenging *C. auris* cells with FK20 RPM (0–200 µg/ml) and caspofungin (0–3.2 µg/ml). The MIC₅₀ values for FK20 RPM and caspofungin in combination were 3.125 µg/ml and 1.05 µg/ml, respectively (Figure 6A). The fractional inhibitory concentration index (FICI), calculated using established methods (57), yielded a value of 0.378, confirming a synergistic interaction between FK20 RPM and caspofungin.

**Figure 6:**
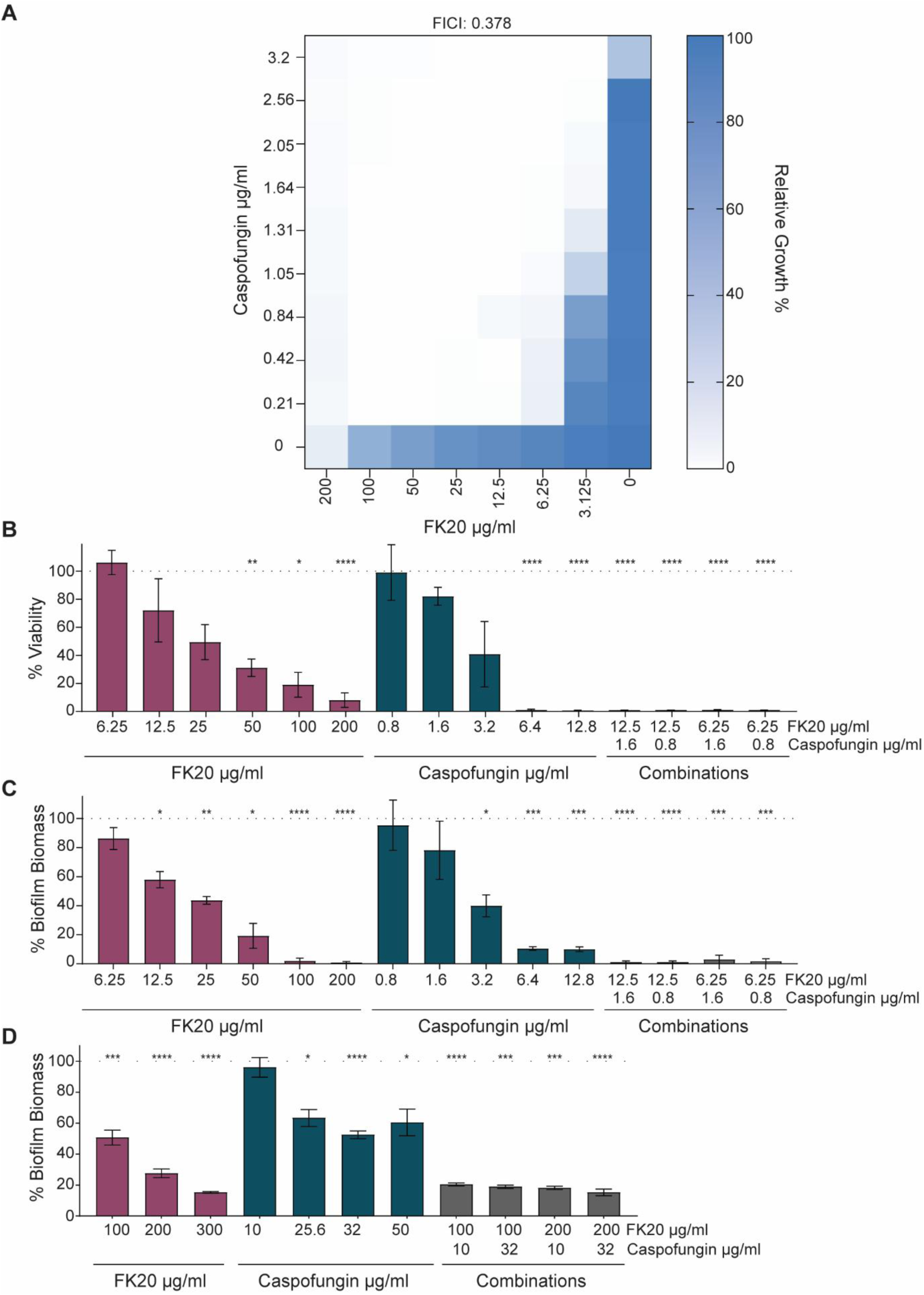
FK20 synergizes with caspofungin to exert fungicidal activity against *C. auris*, inhibit biofilm formation, and disrupt pre-formed mature biofilms. (**A**) Combined effects of FK20 and caspofungin on *C. auris* growth assessed by checkboard assay. Fractional Inhibitory Concentration Index (FICI) values were calculated, and the heat map depicts percentage growth under various combinations of caspofungin and FK20 concentrations. **(B-C)** Inhibition of *C. auris* biofilm formation by FK20, caspofungin, or their combination, quantified after 24 h using the MTT assay (**B**) and crystal violet staining (**C**). FK20 and caspofungin were present in the growth medium from the start of the experiment. **(D)** Destruction of pre-formed mature *C. auris* biofilms following treatment with FK20, caspofungin, or their combination. *C. auris* cells (1 x 10^5^ cells/ml) were incubated in RPMI-1640 supplemented with supplemented with L-Glutamine (1%) and glucose (2%) and grown in a static culture for 24h at 37°C to allow formation of mature biofilm. Planktonic cells were removed and pre-formed, mature biofilms were washed with UPW. FK20, Caspofungin or their combinations were added at the indicated concentrations and incubated at 37C for 24 hours. Biofilm biomass was determined by crystal violet staining. Data represent mean percentage biomass or viability ±SEM of three biologically independent experiments performed with three technical replicates per condition.

Biofilm formation is a critical virulence factor in *Candida* species and plays a central role in *C. auris* persistence, immune evasion, and antifungal tolerance (58). Biofilm-associated phenotypes can be interrogated at distinct stages that reflect different clinical scenarios. Inhibition of biofilm formation models early intervention settings, such as initial host colonization, catheter insertion, or prophylactic antifungal exposure, where preventing adhesion and early biofilm maturation can limit persistence, dissemination, and the development of antifungal tolerance (59,59,60). This is particularly relevant for *C. auris*, which readily colonizes skin and medical devices and serves as a reservoir for nosocomial transmission (61,62). In contrast, assays targeting pre-formed mature biofilms address established infections, which are typically far more tolerant to antifungal therapy due to metabolic dormancy and extracellular matrix–mediated drug exclusion. For instance, caspofungin exhibits limited efficacy against *C. auris* biofilms, requiring concentrations (>32 µg/mL for 90% inhibition) that exceeds safe clinical doses (58,63). Therefore, assessing both biofilm prevention and activity against mature biofilms provides complementary and clinically relevant insights into antifungal efficacy.

Given the synergistic interaction between FK20 RPM and caspofungin, their combined potential to inhibit biofilm formation was evaluated. We first evaluated the ability of FK20 RPM and caspofungin, alone and in combination, to prevent biofilm formation. A biofilm inhibition assay was conducted by culturing *C. auris* cells in a polystyrene 96-well plate with varying concentrations of FK20 RPM, caspofungin, or their combinations. After 24 hours, biofilms were quantified using MTT and crystal violet assays (Figure 6B-C). Individually caspofungin and FK20 reduced biofilm formation in a concentration-dependent manner, achieving complete inhibition of biofilm viability, as determined by MTT assay, at 200 µg/ml FK20 and 6.4 µg/ml caspofungin, and complete inhibition of biofilm biomass as determined by crystal violet staining, at 100 µg/ml FK20 and 12.8 µg/ml caspofungin, respectively. Neither agent inhibited biofilm formation at low concentrations. In contrast, four combinations of FK20 RPM and caspofungin completely inhibited biofilm formation at concentrations 16-fold lower for FK20 and 8-fold lower for caspofungin than those required when each agent was used alone. Bliss independence analysis of biofilm formation inhibition, quantified by crystal violet staining, yielded positive Bliss scores ranging from 0.44 to 0.81, consistent with synergistic interactions, with the highest Bliss score observed at the lowest concentrations of both compounds. These findings highlight the potent synergistic efficacy of FK20 and caspofungin in preventing *C. auris* biofilm formation at clinically relevant concentrations.

Given the marked resistance of established *C. auris* biofilms to antifungal therapy, we next examined the activity of FK20 RPM against pre-formed (24 h) mature biofilms, both alone and in combination with caspofungin (Figure 6D). Consistent with previous reports, caspofungin alone did not significantly reduce mature *C. auris* biofilms biomass at therapeutically relevant concentrations up to 10 µg/ml, with only partial biofilm reduction observed at higher, toxic concentrations (>25 µg/ml). In contrast, FK20 RPM exhibited substantial antibiofilm activity against mature biofilms as a single agent, resulting in dose-dependent reductions in biofilm biomass of approximately 49% at 100 µg/ml, 73% at 200 µg/ml, and 85% at 300 µg/ml.

Notably, the FK20–caspofungin combination displayed synergistic activity against mature biofilms at FK20 concentrations (100 µg/ml) that are two-fold lower than its MIC in planktonic culture, when combined with caspofungin at 10 µg/ml, as determined by Bliss independence analysis (S_Bliss = 0.3). While synergy scores declined at higher drug concentrations, this observation is notable given that synergistic interactions against pre-formed *C. auris* biofilms are rarely observed, even for combinations that are highly synergistic against planktonic cells (64). Together, these data indicate that FK20 RPM retains potent antibiofilm activity against mature *C. auris* biofilms and can sensitize biofilm-embedded cells to caspofungin at otherwise ineffective concentrations.

### FK20 RPM demonstrates therapeutic potential in a murine candidiasis model

To evaluate the therapeutic potential of FK20 RPM, its cytotoxicity was assessed *in vitro* using RAW 264.7 murine macrophages, with IK20 RPM as a control. Cells were incubated with increasing concentrations of each RPM for 24 hours, and viability was measured using the MTT assay (Figure S3). The control 10-mer and 20-mer IK RPMs exhibited no cytotoxicity at concentrations up to 400 µg/ml. In contrast, FK20 RPM induced approximately 13% cytotoxicity at 200 µg/ml, a concentration effective against *C. auris*. At 400 µg/ml, twice its MIC, FK20 RPM exhibited pronounced cytotoxicity (59%) in macrophages, whereas IK20 and IK10 RPMs remained non-cytotoxic. These results indicate a concentration-dependent cytotoxic effect of FK20 RPM and suggest a therapeutic window in which antifungal activity is achieved with limited host-cell toxicity. To confirm FK20’s safety *in vivo*, mice received daily intramuscular injections of FK20 RPM)25 mg/kg), IK20 RPM (25 mg/kg, control), or ultra-pure water (UPW) for four days. The FK20 dosing regimen and route of administration were selected based on prior *in vivo* studies demonstrating efficacy, stability, effective drug distribution, and tolerability of FK20 random peptide mixtures in murine infection models. In these studies, FK20 administration via intramuscular or intravenous injection did not induce hemolysis, abnormal serum chemistry, or histopathological signs of toxicity, and was well tolerated at doses equal to or exceeding those used here (45,65). The present dosing strategy therefore reflects a literature-supported, conservative approach for early *in vivo* evaluation. On day four, blood chemistry and hemolysis were analyzed (Figure S4, Table S1). No significant differences were observed in hemolysis or blood chemistry profiles between FK20-treated and IK10 or UPW-treated cohorts, indicating FK20’s safety for therapeutic use. Although systemic FK20 concentrations were not directly measured in this study, prior work has demonstrated that FK20 random peptide mixtures are stable *in vivo* for at least 24 h following intramuscular or intravenous administration in mice (45,65). This stability indicates resistance to host proteolytic degradation and supports biologically meaningful exposure during the treatment period.

To evaluate the *in vivo* efficacy of FK20 RPM, a neutropenic mouse model of invasive candidiasis was employed, involving intravenous *C. auris* challenge and assessment of treatment response *via* kidney fungal burden (66). Immunosuppression was induced with cyclophosphamide, followed by intravenous *C. auris* infection. Mice received daily intra-muscular injections of FK20 RPM (25 mg/kg), or IK20 RPM (25 mg/kg, control), which lacks antifungal activity. Four days post infection, disease severity, survival, weight loss, and kidney fungal burden were assessed (Figure 7A-E). Compared to IK20-treated mice, FK20-treated mice exhibited significantly reduced clinical scores, increased survival rates, attenuated weight loss, and lower fungal burden. These findings demonstrate that FK20 RPM effectively mitigates acute *C. auris* infection *in vivo*, highlighting its therapeutic potential for systemic candidiasis. In addition to its antifungal efficacy, FK20 treatment was well tolerated *in vivo*. FK20-treated mice did not exhibit significant weight loss, abnormal behavior, or overt clinical signs of toxicity during the course of the experiment. Analysis of serum chemistry and hemolysis on day four revealed no significant differences between FK20-treated and control animals, indicating an absence of detectable acute toxicity at the administered dose. Together, these results demonstrate that FK20 random peptide mixtures exhibit *in vivo* antifungal activity against *C. auris* while remaining well tolerated under the conditions tested.

**Figure 7.**
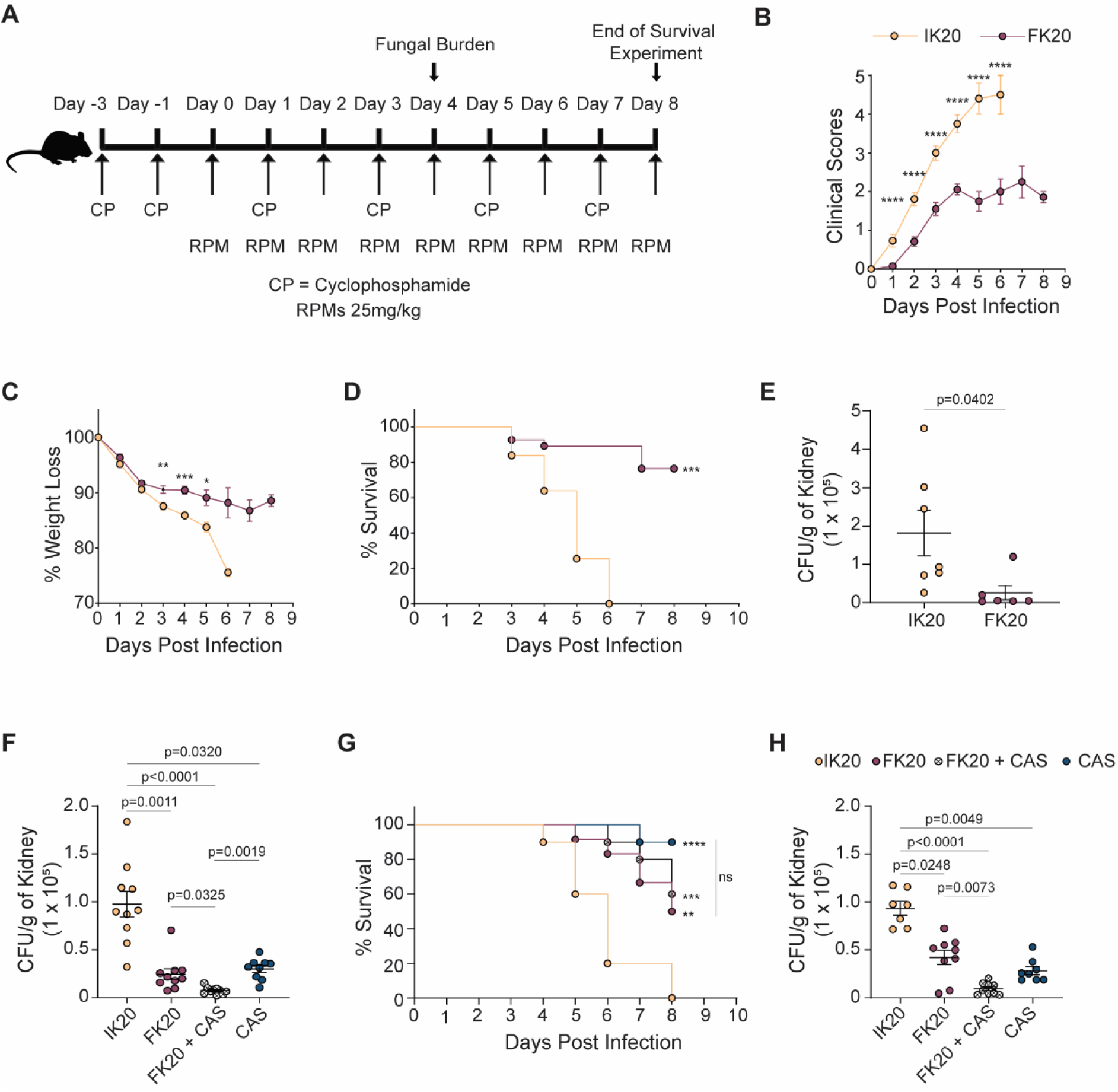
FK20 exhibits potent fungicidal activity in a murine model of systemic candidiasis. (**A**) Schematic overview of the experimental timeline for the murine systemic candidiasis model. C57BL/6 mice were immunosuppressed with cyclophosphamide and subsequently infected intravenously with 1 x 106 CFUs of *C. auris*. Mice received daily intramuscular injections of FK20 or IK20 RPMs (25 mg/kg) throughout the study unless otherwise stated. **(B–C)** Disease severity assessed by semiquantitative clinical scores **(B)** and percentage weight loss **(C)**. Clinical scores (0–5) were assigned based on assessments of fur condition, posture, activity, and lethargy, with higher scores indicating more severe illness. **(D)** Survival percentage of *C. auris*-infected mice treated with 25 mg/kg of FK20 or IK20 RPMs over a period of 8 days post infection (*n* = 10 per group). **(E)** Fungal burden (CFU/g kidney tissue) on day 4 post-infection in mice treated with FK20 or IK20 RPMs. Data are presented as mean ± SEM (*n* = 7 for IK20, *n* = 6 for FK20). **(F)** Fungal burden in kidneys of mice treated with FK20 or IK20 RPMs (25 mg/kg), caspofungin (5 mg/kg), or their combination (25mg/kg FK20 + 5mg/kg caspofungin); *n* = 10 per group. **(G)** Corresponding percentage survival of mice under the same treatment regimens as **(F)** for a period 8 days post infection (*n* = 10 per group). **(H)** Fungal burden in kidneys of mice treated with reduced doses of FK20 or IK20 RPMs (15 mg/kg), caspofungin (3 mg/kg), or their combination; *n* = 10 per group. Data is expressed as CFU/g of kidney tissue. Statistical analysis: (B-C, E) Student’s t-test (p < 0.05); (D,G) log-rank (Mantel–Cox) test; (F,H) one-way ANOVA with Tukey’s multiple comparisons test. Significance levels are indicated as **p < 0.01, ***p < 0.001, and ****p < 0.0001; ns, not significant.

Given the observed *in vitro* synergy and collateral sensitivity between FK20 and caspofungin, we next assessed the efficacy of combination therapy *in vivo* (Figure 7F-H). Neutropenic mice were intravenously infected with *C. auris* and treated daily with FK20 RPM (25 mg/kg), caspofungin (5 mg/kg), or the combination of both agents. At these doses, FK20 and caspofungin monotherapies each resulted in a substantial reduction in kidney fungal burden compared to control-treated mice (approximately 70% reduction). Combination treatment further decreased fungal burden (92% inhibition relative to control); however, this level of inhibition was consistent with the expected additive effect based on the individual drug activities, and Bliss independence analysis did not indicate a synergistic interaction under these conditions. To increase the dynamic range for detecting enhanced combination effects, additional experiments were performed using reduced drug doses (FK20, 15 mg/kg; caspofungin, 3 mg/kg). Under these conditions, FK20 monotherapy exhibited a modest reduction in efficacy (57% reduction compared to IK20-treated control), whereas caspofungin retained strong antifungal activity (approximately 70% reduction compared to IK20-treated control). Combination treatment resulted in 90% inhibition of fungal burden relative to control, closely matching the inhibition predicted from additive drug effects and yielding a Bliss score indicative of additive, rather than synergistic, activity. Notably, the high potency of each monotherapy under both dosing regimens limited the dynamic range available for resolving additive versus synergistic interactions *in vivo*.

Together, these findings establish FK20 as a potent antifungal agent against *C. auris*, exhibiting fungicidal activity *in vitro*, efficacy against biofilm formation and mature biofilms, and therapeutic activity in a murine model of systemic candidiasis. In addition, repeated exposure experiments failed to select for FK20-resistant *C. auris* populations under conditions that readily permitted resistance to conventional antifungals, indicating a high barrier to resistance development. FK20 displays collateral sensitivity with caspofungin and acts synergistically with this agent *in vitro*, resulting in enhanced antifungal activity. *In vivo*, both FK20 and caspofungin monotherapies significantly reduced fungal burden, and combination treatment further improved infection control, achieving near-complete clearance under the conditions tested. Collectively, these results support FK20 as a robust antifungal candidate with durable activity and potential utility both as a standalone therapy and in combination with echinocandins.

## Discussion

The escalating threat of antimicrobial resistance, underscored by reports from the World Health Organization and the CDC, represents a global public health crisis (8,67). While *C. albicans* remains prevalent, the rise of non-*albicans Candida* species, particularly multidrug-resistant *C. auris*, has shifted the epidemiology of candidiasis over the past decade, with *C. auris* classified as an urgent threat by the CDC due to high treatment failure rates (24). Current antifungals, including echinocandins, amphotericin B, and azoles, are limited by narrow spectra, toxicity, pharmacokinetic variability, and drug interactions. Echinocandins, despite favorable safety profiles, are susceptible to resistance in *C. auri*s, while amphotericin B and azoles are constrained by toxicity and drug-drug interactions, respectively (68,69). These limitations, coupled with emerging resistance, necessitate novel antifungal agents with unique mechanisms and robust resistance profiles.

Our study demonstrates that FK20 RPM, a homochiral L-phenylalanine–L-lysine 20-mer, exhibits broad-spectrum antifungal activity with species-specific potency across major human fungal pathogens, including *Candida* spp., *C. neoformans*, and *A. fumigatus*, with particularly high activity against *C. auris*. RPMs are explicitly designed to recapitulate key physicochemical features of AMPs—cationic charge, amphipathicity, and short chain length—while avoiding reliance on a single defined sequence or tertiary structure. Accordingly, FK20’s mode of action is best understood as a collective, stochastic physicochemical attack on fungal membranes rather than a target-specific interaction. Multiple orthogonal readouts support this model: FK20 exposure induced extensive cell surface disruption (SEM), concentration-dependent membrane permeabilization (PI and Ghost Dye uptake), membrane depolarization (DiBAC₄(5)), and oxidative stress (DHR123), all hallmarks of membrane-active antimicrobial agents (Figure 4, Figure S2). Fluorescence microscopy and flow cytometry further demonstrate FK20 association with and internalization into *C. auris* cells (Figure 3); however, this internalization is most parsimoniously interpreted as a consequence of membrane compromise rather than an active uptake mechanism, a distinction well established for AMPs and membrane-disrupting antibiotics (70,71). Consistent with this interpretation, fluorescein-labeled FK20 retained full antifungal activity and showed robust cellular association, whereas fluorescein-labeled control RPMs with minimal antifungal activity (IK10 and LK20) and free fluorescein failed to accumulate in viable fungal cells. Together, these findings support a selective, physicochemically driven interaction with fungal membranes.

The preferential activity of FK20 against fungal cells likely arises from fundamental differences in lipid composition, membrane biophysics, and surface electrostatics between fungal and mammalian membranes, together with the intrinsic physicochemical properties of RPMs. Fungal plasma membranes are enriched in negatively charged phospholipids such as phosphatidylserine and phosphatidylinositol, which are more accessible at the outer leaflet than in mammalian membranes dominated by zwitterionic lipids (72,73). This generates strong electrostatic attraction for cationic, amphipathic peptides like FK20, enabling preferential fungal membrane binding independent of receptors or active uptake (74). Sterol composition further enhances selectivity. Ergosterol-rich fungal membranes differ from cholesterol-containing mammalian membranes in packing, fluidity, and elasticity, rendering them more susceptible to peptide insertion and destabilization, while cholesterol confers resistance to membrane thinning and curvature stress in mammalian cells (75,76). RPMs amplify these intrinsic differences through a collective, stochastic mode of action. Consisting of thousands to millions of short, amphipathic peptides lacking fixed tertiary structure, RPMs insert in multiple orientations and generate distributed nanoscale disruptions rather than discrete pores, preferentially destabilizing membranes with high negative surface potential and lower elastic moduli—features characteristic of fungal membranes.

This framework also explains the observed species-specific differences in susceptibility. FK20 displayed reduced potency against *A. fumigatus* compared with both *Candida* spp. and *C. neoformans* (Fig. 2A), consistent with the exceptionally high melanin content of *A. fumigatus* conidia (∼5–10% of cell wall dry weight) relative to *Candida* spp. and *C. neoformans* (∼1%). Melanin is a negatively charged, hydrophobic polymer that acts as a physical and chemical barrier by sequestering cationic peptides, limiting their access to the plasma membrane, and scavenging reactive oxygen species, thereby attenuating AMP-mediated killing. These properties are well documented to reduce the activity of membrane-active antimicrobial peptides and support a physicochemical, rather than target-specific, mode of action (77,78). In contrast, *C. neoformans* was highly susceptible not only to FK20 but also to IK20 (Fig. 2A), despite the lower hydrophobicity of isoleucine compared with phenylalanine. This heightened sensitivity likely reflects differences in membrane composition, particularly the lower ergosterol content of *C. neoformans*, which may render its plasma membrane more vulnerable to even moderately hydrophobic RPMs. Accordingly, IK20—while minimally fungicidal against *Candida* spp. and *A. fumigatus*—retains appreciable activity against *C. neoformans*, illustrating how subtle changes in RPM hydrophobicity can selectively tune membrane interactions across fungal species.

Rather than representing a limitation, the absence of a specific molecular target constitutes a major therapeutic advantage, particularly against pathogens such as *C. auris* that rapidly acquire target-based drug resistance. Unlike many natural AMPs, which may rely on defined secondary structures or stereospecific interactions, RPMs comprise thousands to millions of related but non-identical sequences. This intrinsic heterogeneity eliminates a conserved binding epitope, preventing fungi from evolving resistance through receptor modification or sequence-specific antagonism. Importantly, this generalized membrane-targeting mechanism imposes a high barrier to resistance evolution. Fungi cannot substantially alter membrane charge, lipid composition, or ergosterol content without incurring severe fitness costs.

Consistent with this principle, a particularly notable finding of this study is the striking asymmetry in resistance evolution and cross-susceptibility between FK20 RPM and caspofungin (Figure 5D-E). Under identical experimental evolution conditions, *C. auris* rapidly acquired high-level resistance to caspofungin, with reduced MIC₅₀, consistent with extensive clinical and experimental evidence demonstrating the low genetic barrier to echinocandin resistance (79–81). In contrast, prolonged exposure to FK20 failed to produce any detectable increase in FK20 MIC, indicating an exceptional inability of *C. auris* to adapt to this compound over multiple passages. Beyond the absence of resistance, caspofungin-resistant lineages exhibited pronounced collateral sensitivity to FK20, whereas FK20-evolved strains retained full susceptibility to caspofungin. The observed collateral sensitivity of caspofungin-resistant *C. auris* strains to FK20 RPM is consistent with known consequences of echinocandin resistance. Resistance-associated alterations in β-1,3-glucan synthesis and compensatory cell wall remodeling are expected to perturb cell wall–membrane coupling and increase membrane vulnerability (82–84), thereby sensitizing cells to membrane-disrupting agents such as FK20. Such unidirectional collateral sensitivity is uncommon among antifungal agents and has important therapeutic implications. The finding that caspofungin resistance enhances susceptibility to FK20 suggests a built-in evolutionary constraint: selection for resistance to one agent actively increases vulnerability to the other. In a combination setting, this creates opposing selective pressures that disfavor the stable emergence of caspofungin resistance, as resistant variants are preferentially eliminated by FK20. Drug pairs exhibiting this resistance–sensitivity trade-off are therefore considered particularly robust, as they both delay resistance evolution and maintain efficacy against resistant subpopulations.

Importantly, combination therapies are predicted to be most effective when such evolutionary constraints coincide with true pharmacological synergy. While collateral sensitivity limits the evolutionary escape routes available to the pathogen, synergy ensures enhanced killing at lower drug concentrations across multiple physiological states. In this context, our finding that FK20 and caspofungin not only exhibit unidirectional collateral sensitivity but also act synergistically, positions this combination as an especially attractive therapeutic pairing (Figure 6). This dual evolutionary–pharmacological advantage provides a strong rationale for the FK20–caspofungin combination and underpins the enhanced activity observed in planktonic cultures, during biofilm formation, and against established biofilms. Mechanistically, this synergy likely reflects complementary modes of action: FK20 disrupts fungal cell walls and membranes through physicochemical interactions, while caspofungin inhibits (1→3)-β-D-glucan synthase, compromising cell wall biosynthesis. Such convergent envelope damage, which exploits fundamental biophysical vulnerabilities rather than single molecular targets, is a recurring theme in successful antimicrobial combination therapies (85,86). Consistent with this model, FK20–caspofungin combinations enabled complete inhibition of *C. auris* biofilm formation at concentrations 16-fold and 8-fold lower than the individual inhibitory concentrations of FK20 and caspofungin, respectively (Figure 6B–C). Biofilm prevention assays capture early stages of fungal adhesion and community establishment, a clinically relevant window in which intervention may avert the development of highly drug-tolerant biofilm structures on host tissues or medical devices. Importantly, biofilm-associated phenotypes represent distinct biological states, with profound implications for antifungal susceptibility. While biofilm prevention assays model early stages of biofilm development, assays using pre-formed biofilms interrogate activity against highly structured, metabolically heterogeneous, and drug-tolerant fungal communities. Mature *C. auris* biofilms are particularly recalcitrant to antifungal therapy, often requiring caspofungin concentrations exceeding clinically achievable levels (>32 µg/mL for ∼90% inhibition), a limitation consistently reported for echinocandins and other antifungal classes (79). In this context, our new data demonstrating that FK20 alone significantly inhibits pre-formed *C. auris* biofilms—and retains moderate but measurable synergy with caspofungin at sub-MIC concentrations—are particularly noteworthy. Unlike many standard antifungals, whose activity is largely restricted to planktonic or early biofilm stages, FK20 exhibits intrinsic antibiofilm activity against established biofilms, a phenotype rarely observed for azoles or echinocandins and more characteristic of certain antimicrobial peptides (58,87,88).

Several features of random peptide mixtures may underlie this activity. The extracellular matrix of fungal biofilms is highly negatively charged, favoring electrostatic accumulation of cationic RPMs (89,90). In addition, the intrinsic sequence diversity of RPMs likely limits proteolytic inactivation, as no single protease can efficiently degrade the entire mixture. RPM-mediated membrane permeabilization and ion imbalance may further enable killing of metabolically dormant biofilm-embedded cells, which are largely tolerant to drugs targeting active biosynthesis (91,92). Finally, resistance mechanisms based on matrix shielding or reduced penetration may be less effective against a heterogeneous, multi-sequence attack that operates through collective physicochemical interactions rather than a single molecular target.

FK20’s therapeutic potential is further supported by its safety and efficacy in an early preclinical, proof-of-concept setting. *In vitro* FK20 exhibited minimal cytotoxicity to RAW 264.7 macrophages at 200 µg/ml —corresponding to twice the MIC required for *C. auris* inhibition—whereas cytotoxic effects were observed only at higher concentrations (400 µg/ml), indicating a therapeutic window between antifungal activity and host cell toxicity (Figure S4). *In vivo*, FK20 (25 mg/kg) showed no adverse effects on blood chemistry or hemolysis in mice (Figure S5, Table S1), addressing common toxic liabilities associated with cationic antimicrobial peptides, including membrane damage and red blood cell lysis, and supporting its overall tolerability at the administered dose. This dosing regimen and route of administration were selected based on prior *in vivo* studies demonstrating that FK20 RPMs are well tolerated in mice, with no evidence of hemolysis, abnormal serum chemistry, or histopathological toxicity following intramuscular or intravenous administration (45,65). Although systemic FK20 concentrations and formal PK/PD relationships were not determined here, FK20 peptide mixtures have previously been shown to be stable *in vivo* for at least 24 h following administration in mice, indicating resistance to host proteolytic degradation and supporting sustained exposure during treatment.

In a neutropenic mouse model of candidiasis, FK20 significantly reduced fungal burden, clinical scores, and weight loss, and increased survival compared to IK20-treated controls (Figure 7B–E), demonstrating *in vivo* antifungal efficacy in a relevant model of systemic infection. These findings are consistent with the potent fungicidal activity observed *in vitro* and further validate FK20 as a promising candidate for the treatment of invasive *Candida* infections.

Given the pronounced *in vitro* synergy and collateral sensitivity observed between FK20 and caspofungin, we next explored the therapeutic potential of combination treatment *in vivo* (Figure 7F-H). At the doses tested, both FK20 and caspofungin monotherapies were already highly effective, each producing substantial reductions in kidney fungal burden. Combination therapy resulted in a further decrease in fungal burden relative to either monotherapy alone; however, this additional reduction was quantitatively consistent with the level of inhibition predicted from additive drug effects, as assessed by Bliss independence analysis. Similar results were obtained when drug doses were reduced to increase the dynamic range for detecting enhanced combination effects, as caspofungin retained strong activity and FK20 remained potently antifungal. Under these conditions, combination treatment again achieved fungal burden reductions closely matching additive expectations, without evidence for *in vivo* synergy. These findings suggest that the high intrinsic potency of each agent in this acute infection model may impose a ceiling effect that limits the ability to discriminate between additive and synergistic interactions *in vivo*, a challenge that has been noted in other systemic antifungal combination studies (93–96). Importantly, the absence of detectable *in vivo* synergy under these conditions does not negate the therapeutic relevance of the FK20–caspofungin combination. Combination therapy consistently achieved near-maximal fungal clearance and may provide advantages in clinical contexts not fully captured by short-term fungal burden measurements, such as mitigation of resistance emergence, enhanced efficacy against heterogeneous fungal populations, or improved outcomes in infections caused by echinocandin-resistant isolates. Indeed, the robust *in vitro* synergy observed across multiple assays, together with the failure to select FK20-resistant *C. auris* under prolonged exposure, supports the rationale for further investigation of FK20-based combination regimens in models explicitly designed to probe resistance suppression and long-term treatment outcomes.

The scope of the *in vivo* evaluation presented here is consistent with prevailing practices in early-stage antimicrobial peptide research. Numerous AMP studies advance candidates into murine infection models based on a combination of *in vitro* cytotoxicity and hemolysis assays, together with limited *in vivo* safety readouts such as serum chemistry or short-term tolerability, without comprehensive pharmacokinetic or dose–exposure analyses (97,98). Within this framework, the current study aligns with accepted standards by demonstrating *in vivo* efficacy while ruling out overt acute toxicity using safety endpoints that are directly relevant to the known liabilities of cationic, membrane-active peptides.

A limitation of the present study is that survival was monitored for up to 8 days post infection, reflecting the use of a high-inoculum intravenous candidiasis model associated with rapid disease progression. Within this ethically approved timeframe, FK20 treatment (alone or in combination) resulted in near-complete fungal clearance by day post infection, as assessed by both CFU enumeration and qPCR-based measurements. These findings provide strong evidence for *in vivo* antifungal efficacy, while longer-term outcome studies and infection models optimized to interrogate combination effects will be of interest in future work.

In conclusion, FK20 RPM represents a mechanistically distinct antifungal strategy against *C. auris*, combining potent cell envelope–directed activity with a markedly reduced propensity for resistance development and favorable interactions with existing antifungals. FK20’s ability to exploit collateral sensitivity in caspofungin-resistant strains, together with its robust synergy with caspofungin in planktonic cells and measurable activity against mature biofilms, underscores its promise as a component of combination therapies aimed at overcoming both intrinsic and acquired antifungal resistance. Unlike conventional target-specific antifungals, FK20 retains activity across multiple fungal physiological states, including drug-tolerant biofilm-associated populations, a feature that is increasingly recognized as critical for durable therapeutic success. Importantly, FK20 demonstrated efficacy *in vivo* without overt toxicity, supporting the translational feasibility of random peptide mixtures as antifungal agents. Taken together, our findings suggest that FK20 and related RPMs may constitute a broadly applicable antimicrobial platform, particularly well-suited for infections characterized by biofilm formation, high tolerance, and rapid resistance emergence. Future studies should focus on defining optimal dosing and formulation strategies, expanding evaluation across diverse fungal pathogens and biofilm architectures, and assessing the performance of FK20-based combinations in advanced preclinical models. Such efforts will be essential to determine whether the unique properties of RPMs can be harnessed to complement or extend the current antifungal armamentarium in the clinic.

## Author contributions

Conceptualization: JA, YB, ZH, and NS

Investigation: JA, YB, MR, HH, AI, ZH, and NS.

Funding acquisition: ZH and NS

Supervision: ZH and NS Writing: JA, YB, ZH, and NS

## Materials and Methods

### Peptide Synthesis

Peptides were synthesized by a standard Fmoc-based solid-phase peptide synthesis (SPPS) on Rink Amide resin (substitution 0.6 mmol/gr), using a peptide synthesizer (Liberty Blue). Amino acids were diluted with dimethylformamide (DMF) to a final concentration of 0.2 M. Before coupling execution, 4 equivalents of amino acids were activated in DMF using 4 equiv. of DIC and 8 equiv. Oxyma. Fmoc deprotection was conducted by 20% piperidine in DMF. At the end of the synthesis, the peptide mixtures were cleaved from the resin by adding a solution containing 95% trifluoroacetic acid (TFA), 2.5% double-distilled water (DDW) and 2.5% triisopropylsilane (TIPS) and stirred for 3 hours. The mixture was then filtered, and the peptides precipitated by the addition of 40 ml cold diethyl ether to the TFA solution and centrifuged. The supernatant was then removed and the peptide pellet dried, dissolved in 20% acetonitrile in DDW, frozen with liquid nitrogen and lyophilized. The synthesis was validated by MALDI-TOF mass spectrometry.

### Cells and Reagents

Mouse macrophage-like RAW 264.7 cells were cultured in DMEM medium supplemented with 10% fetal bovine serum, 1% glutamine, 100 units/ml penicillin/streptomycin and maintained at 37°C in 5% CO_2_. Toxicity assay was done using 3-(4, 5,-dimethyl thiazolyl-2)-2,5-diphenyl tetrazolium bromide assay (MTT).

### *C. auris* Killing Assay

Overnight starter of *C. auris* was prepared in 5ml of YPD media, and grew at 37°C, 200 rpm. The starter was washed three times with PBS. Cells were counted using haemocytometer and resuspended in PBS. 10^6^ cells/ml were incubated with 50 or 200 µg/ml of each RPM separately for 45 min at 37°C, 200 RPM in PBS. The microbial burden was calculated after dilutions and plating each sample on YPD agar plates.

### Scanning Electron Microscopy (SEM) imaging of *C. auris*

Overnight starter culture of *C. auris* was grown in YPD at 37°C with shaking at 200rpm. Cells were harvested and washed three times with PBS, counted with hemocytometer and diluted to a final concentration of 5 x 106 cells/ml. Cell suspensions were incubated with 200ug/ml of each RPM in PBS for 45 minutes at 37°C with shaking at 200rpms. This was followed by additional washes with PBS. *C. auris* cells were resuspended in 4% glutaraldehyde and fixed for 1hour at room temperature. Fixed cells were washed with PBS and 30ul of each sample was placed above and below 0.5mm pre-coated poly-L-lysine glass slides and incubated for 1h at room temperature in a humid chamber to promote cell adhesion.

Samples were dehydrated through a graded ethanol series of increasing concentrations and then transferred to small porous baskets in ethanol to allow cells reach critical point drying (Quorum K850) using liquid CO_2_. Ethanol was replaced by 100% liquid CO_2_ at 5-10 °C and the chamber was heated to 35°C so that at 31°C, liquid CO_2_ passed into the gas phase and the resulting pressure was slowly released.

Dried samples were mounted on metal stubs and sputter-coated with an Au/Pd alloy at a ratio of 80:20 using a Q150T ES coater. Micrographs were acquired on a JEOL JSM-7800F field-emission scanning electron microscope at the Faculty of Agriculture, Food, and Environment, The Hebrew University of Jerusalem, operated in high-vacuum mode at 20kV using a secondary electron detector.

### Mice, Animal Care, and Ethics Statement

All experiments involving the use of mice were approved by and performed according to the Hebrew University of Jerusalem Institutional Animal Care and Use Committee (IACUC) (protocol number MD-21-16494-5). The Hebrew University of Jerusalem is accredited by the NIH and by AAALAC to perform experiments on laboratory animals (NIH approval number: OPRR-A01-5011). Eight-week-old C57BL/6JOlaHsd female mice (18-20g) mice were purchased from Envigo, Israel. Mice were housed under specific pathogen-free conditions in groups of five in individually ventilated cages with unrestricted access to food and water.

### Animals and dosing rationale

The FK20 dosing regimen and route of administration were selected based on previously published *in vivo* studies demonstrating efficacy and tolerability of FK20 random peptide mixtures in murine infection models. In these studies, FK20 administration via intramuscular or intravenous injection did not induce hemolysis, abnormal serum chemistry, or histopathological signs of toxicity (45,65). The dose used in the present study is equal to or lower than doses previously shown to be safe and effective against bacterial pathogens *in vivo*. Animals were monitored daily for body weight, behavior, and clinical signs of distress throughout the experiment.

### Murine Model of Systemic Candidiasis

For toxicity studies, mice (cohorts of five) were injected intramuscularly once daily with FK20 or IK10 (25 µg/g body weight), or with sterile ultrapure water (UPW) as a control. Mice were euthanized on day 5, and blood was collected by tail vein sampling. Blood chemistry analyses were performed by a certified veterinary diagnostic laboratory at the University Veterinary Hospital in Rishon LeZion, Israel. Blood chemistry was assessed in unfasted mice; therefore, glucose measurements may reflect procedural stress and were not used to infer basal metabolic status. To evaluate the hemolytic activity of FK20, 200 µL of blood was transferred to a 96-well plate and centrifuged at 1,000 × g for 10 min at 4 °C. The plasma supernatant was then transferred to a fresh 96-well plate and diluted 1:30 (v/v) in PBS. Hemoglobin release was quantified spectrophotometrically by measuring absorbance at 540 nm. Plasma from control mice treated with 2% Triton X-100 served as a positive hemolysis control. These endpoints were selected to detect common toxic liabilities associated with cationic antimicrobial peptides, including membrane damage, red blood cell lysis, and organ injury.

Mice were immunosuppressed by intra-peritoneal injection of 0.2ml cyclophosphamide in PBS at a concentration of 150 mg/kg three days before infection, followed by 100mg/kg one day before infection, and two days post-infection. Subsequent injection of cyclophosphamide (100mg/kg) were administered at two-day intervals following infection in the survival experiment. Infection inoculum was obtained by sub-culturing *C. auris* on agar plates grown at 37°C for 24h. Single colonies of *C. auris* were resuspended in PBS and adjusted to specified concentrations. Immunosuppressed mice were infected with 1 x 10^6^ CFUs in 0.1 ml of PBS *via* the lateral tail vein injection. Treatment of mice with 25 µg/gr body weight of each RPM was initiated approximately 2 hours after infection and repeated daily throughout the course of the experiment. Following infection, the health status and survival of mice were monitored daily. Clinical scores were assigned to each mouse to evaluate disease severity using disease severity indices obtained from observations of mice activity, grooming, lethargy, and posture. The following parameters were included: fur, coat and posture (normal, 0; fur mildly ruffled, 1; fur strongly ruffled, 2; fur strongly ruffled and hunched posture, 3; mild lethargy 4; severe lethargy, 5). The maximum possible score was 5. Mice were humanely sacrificed when they reached the humane endpoints defined as clinical score of ≥ 4, or ≥20% loss of baseline body weight. The survival curves were analyzed by log-rank (Mantel-Cox) test using GraphPad Prism software version 8.00 (GraphPad Software, USA). Mice were sacrificed and the excised kidneys were weighed followed by homogenization using GentleMACS Dissociator (Miltenyi Biotec). Subsequently, 100 µL of organ homogenate was used in the determination of CFUs following a 1:10000 dilution.

### Determination of MIC

Minimum Inhibitory Concentration (MIC) assays for *C. auris* B11117 strains were performed for FK20 and caspofungin according to European Committee on Antibiotic Susceptibility Testing (EUCAST) guidelines for determination of MIC using broth microdilution method (58), (59). Single colonies were obtained for *Candida auris* B1117 strains streaked on Sabouraud Dextrose Agar (40 gr/L Dextrose, 10 gr/L Peptone, at pH 5.6) plates at 37°C for 24h. Approximately 1 x 10^5^ *C. auris* cells were incubated in 200ul filter sterilized RPMI-1640 culture media (Sigma), supplemented with 1.8% glucose to reach a final concentration of 2% and 0.03% L-Glutamine at pH 7.0. The growth media was supplemented with serial dilutions of antifungal agents 0 to 400 µg/ml of FK20 RPM, and 0 to 64 µg/ml of caspofungin in different 96-well microtiter plates. Cells were incubated in 96-well microtiter plates at 37°C and MIC was determined spectrophotometrically after 28h, 48h, and 72h using Tecan Infinite M Plex plate reader (Neotec Bio, Israel).

### Drug Combination Assay and Determination of Fractional Inhibitory Concentration

The combined efficacy of Caspofungin and FK20 was determined *in vitro* using a checkerboard assay, followed by the calculation of the Fractional Inhibitory Concentration Index (FICI). Caspofungin and FK20 were added individually at a volume of 50 µL per well in four-fold strength, yielding a final concentration of 0-3.2 µg/ml for Caspofungin and 0-2000 to 200 µg/ml for FK20. The yeast inoculum (1 x 10^5^ cells per well) was added to the plates. Plates were then incubated at 37°C for a duration of 72 hours. The antifungal combination activities of caspofungin and FK20 were assessed based on their FICI. The FICI was derived from the Minimum Inhibitory Concentration (MIC-50), representing the inhibitory concentration at which 50% of the cell population are killed by the effect of the drug combination. The FICI was determined by the formula; 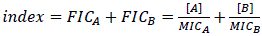 where [A] and [B] are concentrations of drugs A and B, and MIC_A_ and MIC_B_ are their respective MIC values for *C. auris*. Results were interpreted as follows: Synergy (FICI ≤ 0.5), partial synergy (0.5 < FICI ≤ 0.75), additive (0.75 < FICI ≤ 1), indifferent (1 < FICI ≤ 4), and antagonistic (FICI > 4).

### Biofilm Formation and Quantification

In a 96-well microtiter plate, 200 µL of *C. auris* cells at a concentration of 10^6^ CFU/ ml were cultured in RPMI-1640 medium, supplemented with L-Glutamine (1%) and glucose (2%) for 24 hours at 37°C together with different concentrations of a FK20, caspofungin, or combinations that were tested to study their effects on biofilm formation. Inhibition of biofilm formation was assessed using a crystal violet assay and MTT assay. Planktonic cells were removed; biofilms were washed with UPW and then stained with 125 µL of 0.1% crystal violet for 15 minutes. Excess stain was washed off, and the biofilms were air-dried for 4 hours at 25°C. Finally, 125 µL of 30% acetic acid was added to each well and biofilm biomass was quantified at 595 nm using a microplate reader (Tecan infinite M200).

Biofilm viability was assessed using the 3-(4,5-dimethylthiazol-2)-2,5-diphenyltetrazolium bromide (MTT) assay. In this method, each well was loaded with 50 µL of MTT reagent (1.5 mg/ml), and the plate was incubated for 1.5 hours at 37°C. Subsequently, 150 µL of dimethyl sulfoxide (DMSO) was added, and the plate was incubated for 10 minutes at 30°C with agitation at 180 RPM. The absorbance was then measured at 595 nm using a TECAN Infinite plate reader.

To quantify the ability of FK20 to disrupt pre-formed mature biofilms, *C. auris* cells (1 x 10^5^ cells/ml) were grown in RPMI-1640 medium, supplemented with L-Glutamine (1%) and glucose (2%) for 24 hours at 37°C to allow formation of mature biofilms. Planktonic cells were discarded and mature biofilms were washed with PBS. FK20, Caspofungin, and their combinations at specified concentrations were added to each well and incubated at 37°C for 24 hours. Each well was washed twice with UPW, and biofilm biomass was determined using crystal violet assay as described above.

### *In vitro* experimental evolution

*In vitro* experimental evolution of *Candida auris* was performed according to the protocol published in (45) with minor modifications. Briefly, *C. auris* B11117 was independently passaged through 100 µg/ml of FK20 RPM and 2 µg/ml of caspofungin. *C. auris* was also passaged simultaneously without exposure to any antifungal drug. The concentration of FK20 RPM selected for the experimental evolution was equivalent to the predetermined MIC_50_ of the parental *C. auris* B11117 strain. The experimental evolution was initiated by generating three independent replicates of *C. auris* B11117 as the parental strain. Cultures were struck from frozen parental stocks onto Sabouraud Dextrose Agar (40 gr/L Dextrose, 10 gr/L Peptone, at pH 5.6) plates and incubated at 37°C overnight. Each parental replicate was generated from randomly selected single colonies. Subcultures were made in YPD liquid media and incubated at 37°C overnight at 100 rpm. The overnight culture was diluted 1:1000 in RPMI-1640 (supplemented with 1.8% glucose to reach a final concentration of 2% glucose and 0.03% L-Glutamine at pH 7.0) to a final volume of 1ml. RPMI-1640 media was supplemented with 100µg/ml of FK20 and 2.5 µg/ml of caspofungin for replicates evolved with FK20 and caspofungin respectively, and supplemented RPMI-1640 media for replicates evolved without drugs in 96-deep well plates and incubated at 37°C for 72 hr. From cultures of each replicate, 100 µL of each replicate was frozen in 25% glycerol as parental replicates at time zero. Precautions were taken to minimize contaminations while keeping environmental conditions similar. Plates were sealed with gas-permeable membranes to allow gas exchange and maximum growth. The 96-deep well plate was incubated at 37°C without shaking to mimic static growth used in clinical resistance assays. Plates were incubated in plastic trays lined with wet paper towels to minimize evaporation. Following incubation for 72 hr, individual wells were mixed by pipetting and diluted 1:10 into fresh RPMI-1640 supplemented with or without drugs. In total 10 transfers were carried out, and 100 µL of evolved replicate cultures were frozen in duplicates in 25% glycerol for each transfer until after the tenth transfer and maintained at –80°C.

### Determination of Resistance for Evolved Strains

Initial susceptibility of all parental strains was tested using broth microdilution liquid assays to determine the MIC50, that is, the concentration of antifungal agent at which 50 percent of the cell population is eliminated. The liquid assay experiment was carried out according to cell dilution regulations of the European Committee on Antibiotic Susceptibility Testing (EUCAST) guidelines with some modifications (58), (59). Briefly, RPMI incubated at 37°C was used at the base media, and growth was determined as absorbance of cell densities at OD_590_ at 24, 48, and 72 hr post-inoculation. MIC_50_ was numerically calculated as the concentration in which a 50% decrease in turbidity was determined spectrophotometrically.

### Viability Staining

To assess the fungicidal activity of FK20, approximately 1 x 10^6^ cells/ml were incubated with varying concentrations of FK20, ranging from 0 to 400 µg/ml for 45 minutes at 37°C with continuous shaking. Subsequently, treated cells were washed with PBS and resuspended with either Propidium Iodide (1:25 dilution of 0.5mg/ml, BioLegend) or Ghost-Dye (1:500 dilution, Tonbo Biosciences) viability stain for 20 minutes. Residual viability dyes were washed with PBS, and cells were resuspended in PBS for analysis of fluorescence by flow cytometer.

### Reactive oxygen species (ROS) detection

Approximately 1 x 10^6^ cells/ml were incubated with 200 µg/ml of FK20 for 45 minutes at 37°C with continuous shaking at 200 RPMs. Treated and untreated cells were washed with PBS and resuspended with 10µM dihydrorhodamine 123 (DHR123, Cayman Chemical) to detect intracellular ROS by staining for 20 minutes. Residual dyes were washed with PBS, and cells were resuspended in PBS for analysis of fluorescence by flow cytometer.

### Cell membrane depolarization

To test the effect of RPMs on cell membrane depolarization of *C. auris*, 1 x 10^6^ cells/ml were incubated with 200 µg/ml of FK20 for 45 minutes at 37°C with continuous shaking at 200 RPMs. Treated and untreated cells were washed with PBS and resuspended with 5µM bis-(1,3-dibutylbarbituric Acid) pentamethine oxonol (DIBAC₄(5), Santa Cruz) to detect intracellular ROS by staining for 20 minutes. ells were resuspended in PBS for analysis of fluorescence by flow cytometer.

### Statistics and reproducibility

All statistical analyses were performed using GraphPad Prism v.8.0.2. No statistical methods were used to pre-determine sample size. Sample sizes were based on previous publications that achieved statistically significant results (P < 0.05). Unless otherwise noted, all statistical analyses were performed with at least three biologically independent samples. All images are representative of a minimum of three biologically independent samples that represent a minimum of three independent experiments unless otherwise noted. For comparisons between two groups, two-tailed unpaired t-tests were performed.

## Supplementary Data

**Figure S1:**
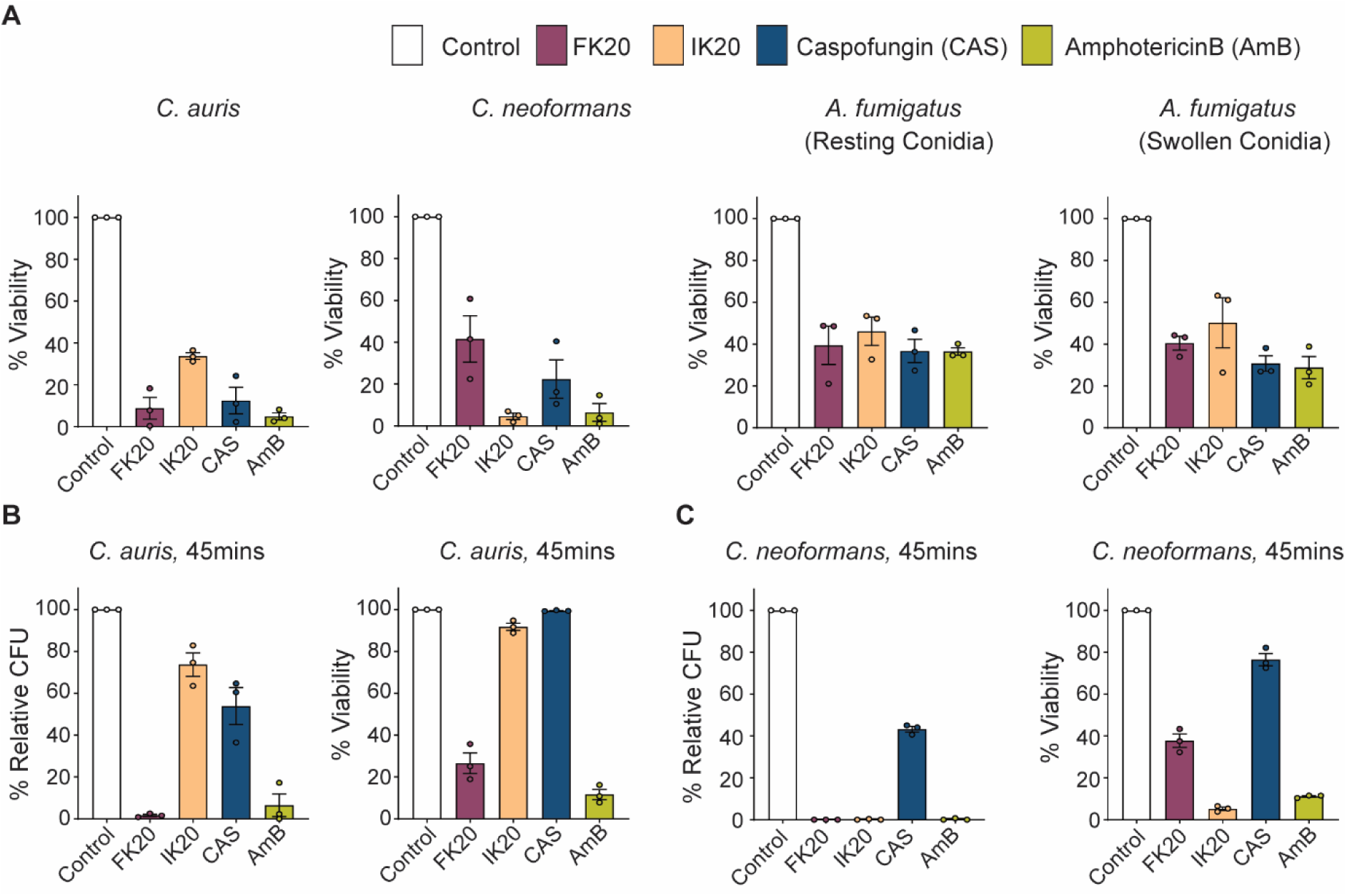
FK20 potency varies across major fungal pathogens of clinical relevance. **A.** *C. auris* and *C. neoformans* yeast cells, as well as *A. fumigatus* conidia (resting and swollen), were incubated with 200µg/ml FK20 or IK20 RPMs, 2µg/ml Caspofungin, 2µg/ml Amphotericin B, or PBS as a control for 5h at 37°C, followed by PI staining to determine cell viability **B-C**. *C. auris* and *C. neoformans* cells were treated as stated above for 45 minutes at 37°C. Fungal viability after treatment is presented for *C. auris* as percentage relative CFUs (B, left panel), % viability after PI staining (B, right panel) and for *C. neoformans* as percentage relative CFUs (C, left panel), % viability after PI staining (C, right panel). Data represents the mean ± SEM of three biologically independent experiments. Statistical significance was determined using student’s t-test, comparing each treatment to the untreated control (set to 100% for each biological replicate). Significance levels are indicated as *p <0.05, **p < 0.01, ***p<0.001 and ****p < 0.0001; values without symbols are not significantly different from the control.

**Supplementary Movie 1. FK20 RPM Penetrates C. auris**. 5·10^7^ CFU/mL of *C. auris* cells were incubated with 200 μg/mL fluorescein-labeled FK20 RPM (green) for 45 minutes at 37°C in PBS. Samples were co-stained with Calcofluor White stain (blue). Z-stack movie prepared using Z images taken by a confocal microscope at a magnification of 40×. The video was edited using ImageJ. Scale bar, 10 μm.

**Figure S2:**
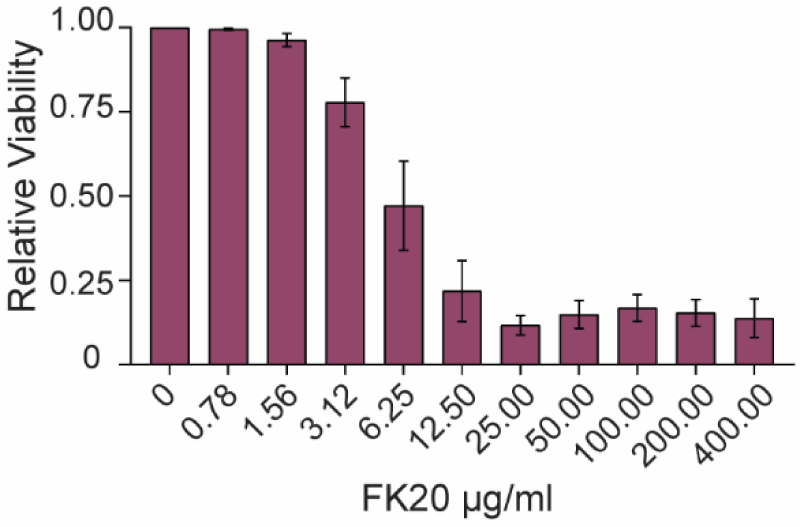
Membrane damage quantification by ghost dye staining. *C. auris* cells were incubated with FK20 RPM at concentration ranging from 0 – 400 µg/mL for 45 minutes at 37°C with continuous shaking. Membrane damage was evaluated by staining *C. auris* cells with Ghost dye for 20 minutes. Cell viability was determined by measuring Ghost dye fluorescence using a flow cytometer. The viability of *C. auris* was calculated as the percentage of cells that remained unstained by Ghost dye relative to the untreated control. The data are derived from three biologically independent experiments conducted in triplicates.

**Figure S3.**
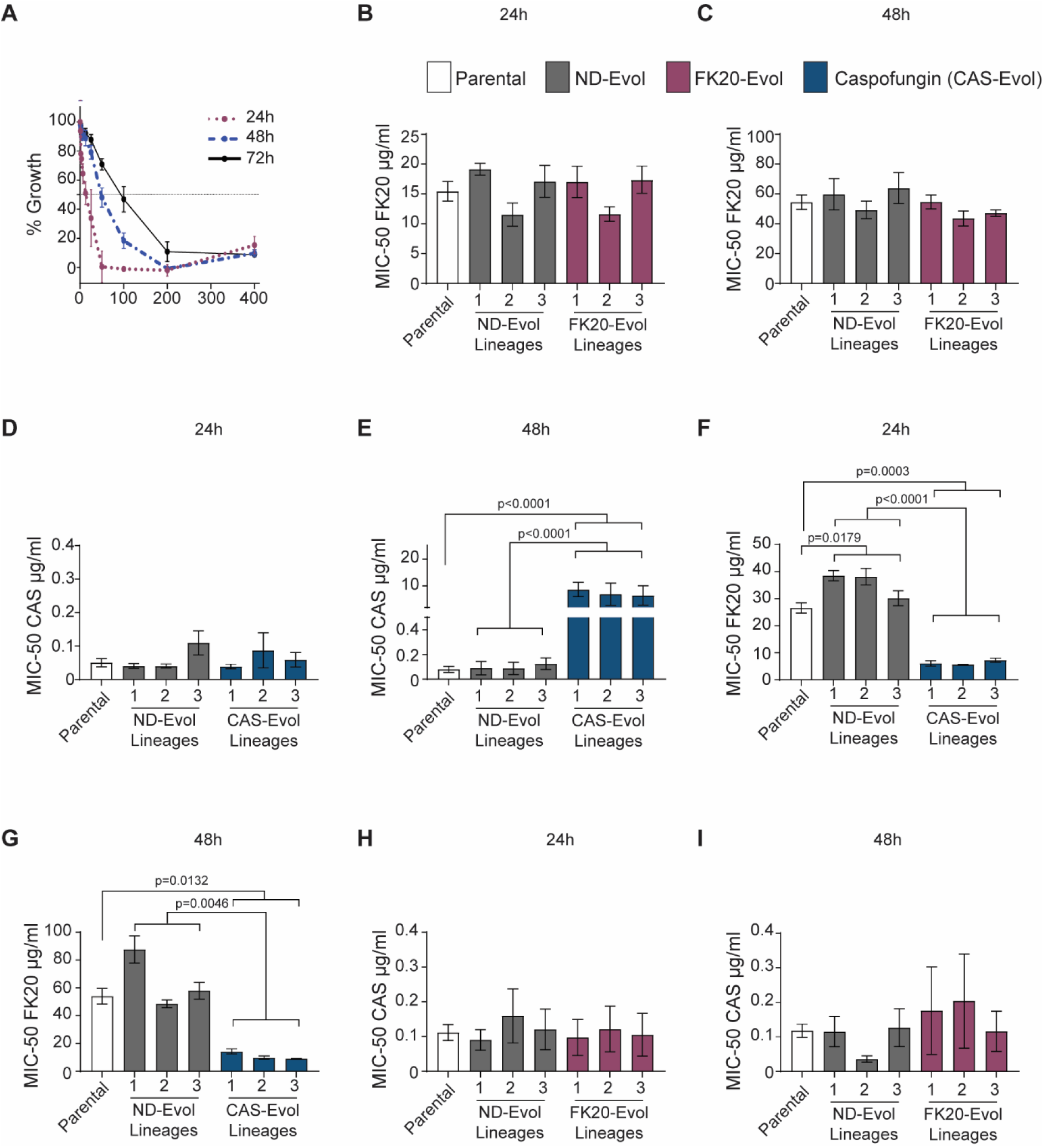
MIC determination and experimental evolution of *C. auris* strains at multiple time points. (**A**) Initial MIC determination for FK20 against parental *C. auris* using broth microdilution assay. Growth curves were measured across FK20 concentrations (0–400 µg/mL) after 24, 48 and 72 hours. **(B–I)** MIC-50 values for parental, non-drug-evolved (ND-evol), and drug-evolved lineages under FK20 or caspofungin treatment at 24 hours **(B, D, F, H)** and 48 hours **(C, E, G, I)** post-exposure. (**B–C)**: FK20 MIC-50 for parental, ND-evol, and FK20-evolved lineages. **D–E**: Caspofungin MIC-50 for parental, ND-evol, and caspofungin-evolved lineages. (**F–G)**: FK20 MIC-50 for parental, ND-evol, and caspofungin-evolved lineages (collateral sensitivity). (**H–I)**: Caspofungin MIC-50 for parental, ND-evol, and FK20-evolved lineages (collateral sensitivity). Each data point represents the mean ± S.E.M. for the parental strains and the three independent lineages of evolved strains across three biological replicates. Statistical comparisons were performed using one-way ANOVA with Tukey’s multiple comparisons test. Significance levels are indicated as *p<0.05, **p<0.01, ***p<0.001 and ****p<0.0001; values without symbols are not significantly different from the control.

**Figure S4:**
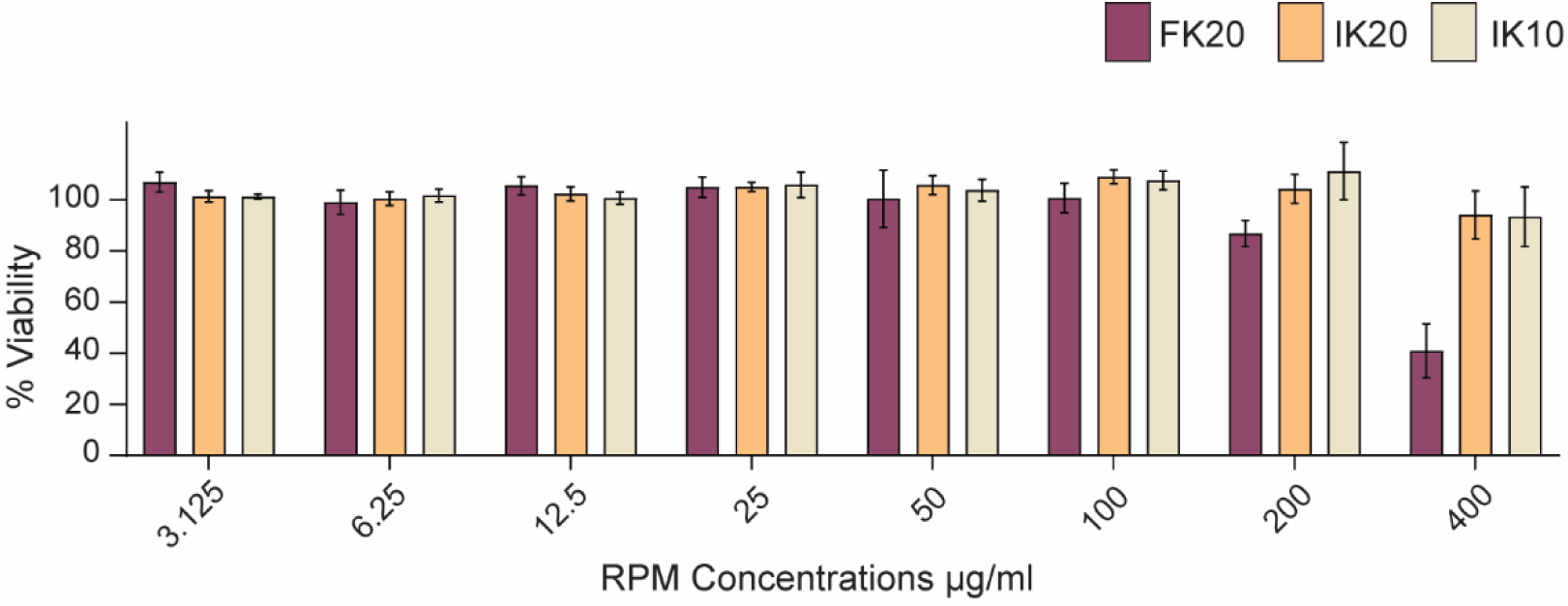
RPMs exhibit no cytotoxicity in cultured cells. RAW 264.7 murine macrophage cells were incubated with FK20, IK20, or IK10 for 24 hours at 37°C in a 5% CO₂ environment. Cell viability was assessed using the MTT assay, with macrophage viability expressed relative to a control group with no treatment. The data represents the mean of three biologically independent experiments, each performed in triplicate.

**Table S1:** Mice blood chemistry. A control group of Mice were administered ultra-pure water, FK20 or IK10 RPMs intramuscularly daily for four days (25 µg/gr body weight). Blood chemistry was analyzed on day four. n=5 per group.

| Test | UPW | IK10 | FK20 | Reference values |
| --- | --- | --- | --- | --- |
| AST (IU/L) | 341.5 ± 246.5 | 187 ± 81 | 275 ± 90 | 57-329 |
| ALT (IU/L) | 38 ± 20 | 28 ± 3 | 32 ± 6 | 22-133 |
| ALP (IU/L) | 141.5 ± 13.5 | 110.5 ± 12.5 | 94 ± 12 | 16-200 |
| GGT (IU/L) | < 3 | < 3 | < 3 | < 3 |
| Triglyceride (mg/dL) | 70 ± 37 | 54.5 ± 20.5 | 61 ± 10 | 16-164 |
| Cholesterol (mg/dL) | 84.15 ± 5.55 | 97.45 ± 3.75 | 85.5 ± 4 | 34-173 |
| Total Bilirubin (mg/dL) | 0.159 | < 0.15 | < 0.15 | 0.1-0.9 |
| Glucose (mg/dL) | 395 ± 178 | 219.5 ± 49.5 | 328 ± 34 | 60-133 |
| Albumin (g/dL) | 3.75 ± 0.55 | 3.8 ± 0.49 | 3.7 ± 0.1 | 2.6-5.4 |
| Total protein (g/dL) | 4.895 ± 0.095 | 4.71 ± 0.44 | 4.6 ± 0.1 | 4.6-7.3 |
| UREA (mg/dL) | 31.4 ± 0.9 | 35.9 ± 0.5 | 32 ± 3 | 2-71 |
| Creatinine (mg/dL) | 0.315 ± 0.015 | 0.32 ± 0.03 | 0.33 ± 0.03 | 0.1-1.8 |
| Phosphate (mg/dL) | 10.85 ± 3.85 | 8.875 ± 0.435 | 9.796 ± 0.6 | 5.3-11.3 |
| Ca (mg/dL) | 9 ± 0.9 | 9.25 ± 0.55 | 8.73 ± 0.33 | 6.8-11.9 |
| Na (mmol/l) | 145 | 149.5 ± 0.5 | 142 ± 1.2 | 145–155 |
| **Cl (mmol/l)** | 104.85 ± 1.25 | 110.55 ± 0.15 | 103 ± 1.36 | 105–115 |
| **K (mmol/l)** | 6.885 ± 0.98 | 6.385 ± 0.135 | 6.5 ± 0.46 | 6.5–9.7 |

**Figure S5:**
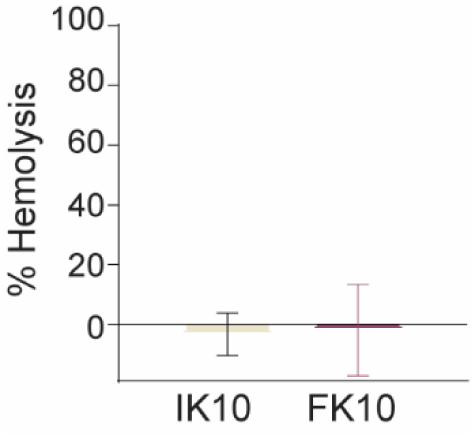
RPMs are not hemolytic. A control group of Mice were administered ultra-pure water, FK20 or IK10 RPMs intramuscularly daily for four days (25 µg/gr body weight). Hemolysis was analyzed on day four by direct spectrophotometric measurement at 540 nm. n=3 per groups.

**Figure S6:**
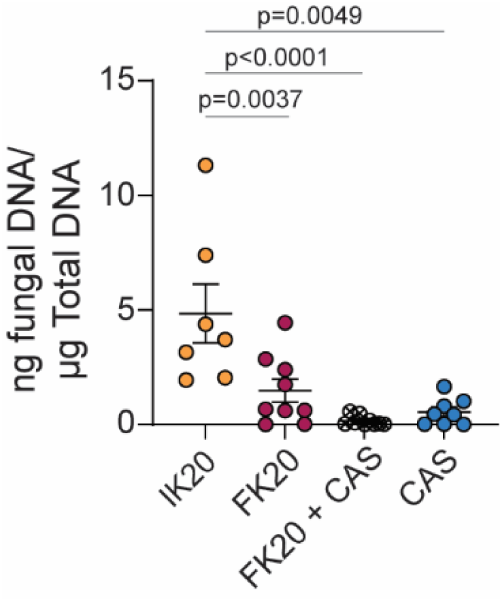
Quantification of fungal burden by qPCR in the murine systemic candidiasis model. Fungal burden was assessed by quantitative PCR (qPCR) on DNA extracted from kidney tissues of *C. auris* infected mice following treatment with IK20 (15mg/kg), FK20(15mg/kg/), Caspofungin (3mg/kg), or a combination of FK20 and Caspofungin (15mg/kg and 3mg/kg respectively). Fungal load was quantified on day 4 post-infection as ng of fungal DNA per µg of total DNA. Statistical analysis was performed using one-way ANOVA, and p-values for significant comparisons are indicated on the graph, comparisons without p-values are not statistically significant.

## References

1. Chinemerem Nwobodo D, Ugwu MC, Oliseloke Anie C, Al-Ouqaili MTS, Chinedu Ikem J, Victor Chigozie U, et al. Antibiotic resistance: The challenges and some emerging strategies for tackling a global menace. Clinical Laboratory Analysis. 2022 Sept;36(9):e24655.

2. Murray CJL, Ikuta KS, Sharara F, Swetschinski L, Robles Aguilar G, Gray A, et al. Global burden of bacterial antimicrobial resistance in 2019: a systematic analysis. The Lancet. 2022 Feb;399(10325):629–55.

3. Nelson RE, Hatfield KM, Wolford H, Samore MH, Scott RD, Reddy SC, et al. National Estimates of Healthcare Costs Associated With Multidrug-Resistant Bacterial Infections Among Hospitalized Patients in the United States. Clinical Infectious Diseases. 2021 Jan 29;72(Supplement_1):S17–26.

4. Zhang Z, Zhang Q, Wang T, Xu N, Lu T, Hong W, et al. Assessment of global health risk of antibiotic resistance genes. Nat Commun. 2022 Mar 23;13(1):1553.

5. Van Rhijn N, Arikan-Akdagli S, Beardsley J, Bongomin F, Chakrabarti A, Chen SCA, et al. Beyond bacteria: the growing threat of antifungal resistance. The Lancet. 2024 Sept;404(10457):1017–8.

6. Horton MV, Holt AM, Nett JE. Mechanisms of pathogenicity for the emerging fungus Candida auris. LeibundGut-Landmann S, editor. PLoS Pathog. 2023 Dec 21;19(12):e1011843.

7. Cristina ML, Spagnolo AM, Sartini M, Carbone A, Oliva M, Schinca E, et al. An Overview on Candida auris in Healthcare Settings. JoF. 2023 Sept 8;9(9):913.

8. Centers for Disease Control and Prevention (U.S.). Antibiotic resistance threats in the United States, 2019 [Internet]. Centers for Disease Control and Prevention (U.S.); 2019 Nov [cited 2025 Dec 23]. Available from: https://stacks.cdc.gov/view/cdc/82532

9. WHO Fungal Priority Pathogens List to Guide Research, Development and Public Health Action. 1st ed. Geneva: World Health Organization; 2022. 1 p.

10. Satoh K, Makimura K, Hasumi Y, Nishiyama Y, Uchida K, Yamaguchi H. *Candida auris* sp. nov., a novel ascomycetous yeast isolated from the external ear canal of an inpatient in a Japanese hospital. Microbiology and Immunology. 2009 Jan;53(1):41–4.

11. Chow NA, Muñoz JF, Gade L, Berkow EL, Li X, Welsh RM, et al. Tracing the Evolutionary History and Global Expansion of Candida auris Using Population Genomic Analyses. Butler G, Nielsen K, editors. mBio. 2020 Apr 28;11(2):e03364–19.

12. Suphavilai C, Ko KKK, Lim KM, Tan MG, Boonsimma P, Chu JJK, et al. Detection and characterisation of a sixth Candida auris clade in Singapore: a genomic and phenotypic study. The Lancet Microbe. 2024 Sept;5(9):100878.

13. Spruijtenburg B, Badali H, Abastabar M, Mirhendi H, Khodavaisy S, Sharifisooraki J, et al. Confirmation of fifth *Candida auris* clade by whole genome sequencing. Emerging Microbes & Infections. 2022 Dec 31;11(1):2405–11.

14. Rhodes J, Fisher MC. Global epidemiology of emerging Candida auris. Current Opinion in Microbiology. 2019 Dec;52:84–9.

15. Welsh RM, Bentz ML, Shams A, Houston H, Lyons A, Rose LJ, et al. Survival, Persistence, and Isolation of the Emerging Multidrug-Resistant Pathogenic Yeast Candida auris on a Plastic Health Care Surface. Diekema DJ, editor. J Clin Microbiol. 2017 Oct;55(10):2996–3005.

16. Tsay S, Kallen A, Jackson BR, Chiller TM, Vallabhaneni S. Approach to the Investigation and Management of Patients With Candida auris, an Emerging Multidrug-Resistant Yeast. Clinical Infectious Diseases. 2018 Jan 6;66(2):306–11.

17. Jangir P, Kalra S, Tanwar S, Bari VK. Azole resistance in *Candida auris*: mechanisms and combinatorial therapy. APMIS. 2023 Aug;131(8):442–62.

18. Chen J, Tian S, Han X, Chu Y, Wang Q, Zhou B, et al. Is the superbug fungus really so scary? A systematic review and meta-analysis of global epidemiology and mortality of Candida auris. BMC Infect Dis. 2020 Dec;20(1):827.

19. Tu J, Zhu T, Wang Q, Yang W, Huang Y, Xu D, et al. Discovery of a new chemical scaffold for the treatment of superbug *Candida auris* infections. Emerging Microbes & Infections. 2023 Dec 31;12(1):2208687.

20. Lockhart SR, Etienne KA, Vallabhaneni S, Farooqi J, Chowdhary A, Govender NP, et al. Simultaneous Emergence of Multidrug-Resistant *Candida auris* on 3 Continents Confirmed by Whole-Genome Sequencing and Epidemiological Analyses. CLINID. 2017 Jan 15;64(2):134–40.

21. Chowdhary A, Prakash A, Sharma C, Kordalewska M, Kumar A, Sarma S, et al. A multicentre study of antifungal susceptibility patterns among 350 Candida auris isolates (2009–17) in India: role of the ERG11 and FKS1 genes in azole and echinocandin resistance. Journal of Antimicrobial Chemotherapy. 2018 Apr 1;73(4):891–9.

22. Chaabane F, Graf A, Jequier L, Coste AT. Review on Antifungal Resistance Mechanisms in the Emerging Pathogen Candida auris. Front Microbiol. 2019 Nov 29;10:2788.

23. Sanyaolu A, Okorie C, Marinkovic A, Abbasi AF, Prakash S, Mangat J, et al. *Candida auris*: An Overview of the Emerging Drug-Resistant Fungal Infection. Infect Chemother. 2022;54(2):236.

24. Chowdhary A, Voss A, Meis JF. Multidrug-resistant Candida auris: ‘new kid on the block’ in hospital-associated infections? Journal of Hospital Infection. 2016 Nov;94(3):209–12.

25. Hu S, Zhu F, Jiang W, Wang Y, Quan Y, Zhang G, et al. Retrospective Analysis of the Clinical Characteristics of Candida auris Infection Worldwide From 2009 to 2020. Front Microbiol. 2021 May 20;12:658329.

26. Geremia N, Brugnaro P, Solinas M, Scarparo C, Panese S. Candida auris as an Emergent Public Health Problem: A Current Update on European Outbreaks and Cases. Healthcare. 2023 Feb 2;11(3):425.

27. Chowdhary A, Anil Kumar V, Sharma C, Prakash A, Agarwal K, Babu R, et al. Multidrug-resistant endemic clonal strain of Candida auris in India. Eur J Clin Microbiol Infect Dis. 2014 June;33(6):919–26.

28. Arendrup MC, Prakash A, Meletiadis J, Sharma C, Chowdhary A. Comparison of EUCAST and CLSI Reference Microdilution MICs of Eight Antifungal Compounds for Candida auris and Associated Tentative Epidemiological Cutoff Values. Antimicrob Agents Chemother. 2017 June;61(6):e00485–17.

29. Huan Y, Kong Q, Mou H, Yi H. Antimicrobial Peptides: Classification, Design, Application and Research Progress in Multiple Fields. Front Microbiol. 2020 Oct 16;11:582779.

30. Pasupuleti M, Schmidtchen A, Malmsten M. Antimicrobial peptides: key components of the innate immune system. Critical Reviews in Biotechnology. 2012 June;32(2):143–71.

31. Raj PA, Dentino AR. Current status of defensins and their role in innate and adaptive immunity. FEMS Microbiology Letters. 2002 Jan;206(1):9–18.

32. Mookherjee N, Anderson MA, Haagsman HP, Davidson DJ. Antimicrobial host defence peptides: functions and clinical potential. Nat Rev Drug Discov. 2020 May;19(5):311–32.

33. Reddick LE, Alto NM. Bacteria Fighting Back: How Pathogens Target and Subvert the Host Innate Immune System. Molecular Cell. 2014 Apr;54(2):321–8.

34. Ageitos JM, Sánchez-Pérez A, Calo-Mata P, Villa TG. Antimicrobial peptides (AMPs): Ancient compounds that represent novel weapons in the fight against bacteria. Biochemical Pharmacology. 2017 June;133:117–38.

35. Kumar P, Kizhakkedathu J, Straus S. Antimicrobial Peptides: Diversity, Mechanism of Action and Strategies to Improve the Activity and Biocompatibility In Vivo. Biomolecules. 2018 Jan 19;8(1):4.

36. Buda De Cesare G, Cristy SA, Garsin DA, Lorenz MC. Antimicrobial Peptides: a New Frontier in Antifungal Therapy. Andrew Alspaugh J, editor. mBio. 2020 Dec 22;11(6):e02123–20.

37. Perez-Rodriguez A, Eraso E, Quindós G, Mateo E. Antimicrobial Peptides with Anti-Candida Activity. IJMS. 2022 Aug 17;23(16):9264.

38. Wang Y, Wang S, Chen Y, Xie C, Xu H, Lin Y, et al. The role of Npt1 in regulating antifungal protein activity in filamentous fungi. Nat Commun. 2025 Mar 23;16(1):2850.

39. Zasloff M. Antimicrobial peptides of multicellular organisms. Nature. 2002 Jan;415(6870):389–95.

40. Hancock REW, Sahl HG. Antimicrobial and host-defense peptides as new anti-infective therapeutic strategies. Nat Biotechnol. 2006 Dec;24(12):1551–7.

41. Fjell CD, Hiss JA, Hancock REW, Schneider G. Designing antimicrobial peptides: form follows function. Nat Rev Drug Discov. 2012 Jan;11(1):37–51.

42. Amso Z, Hayouka Z. Antimicrobial random peptide cocktails: a new approach to fight pathogenic bacteria. Chem Commun. 2019;55(14):2007–14.

43. Topman S, Tamir-Ariel D, Bochnic-Tamir H, Stern Bauer T, Shafir S, Burdman S, et al. Random peptide mixtures as new crop protection agents. Microbial Biotechnology. 2018 Nov;11(6):1027–36.

44. Cheriker H, Stern Bauer T, Oren Y, Nir S, Hayouka Z. Immobilized random peptide mixtures exhibit broad antimicrobial activity with high selectivity. Chem Commun. 2020;56(75):11022–5.

45. Bennett RC, Oh MW, Kuo SH, Belo Y, Maron B, Malach E, et al. Random Peptide Mixtures as Safe and Effective Antimicrobials against *Pseudomonas aeruginosa* and MRSA in Mouse Models of Bacteremia and Pneumonia. ACS Infect Dis. 2021 Mar 12;7(3):672–80.

46. Lau JZ, Kuo SH, Belo Y, Malach E, Maron B, Caraway HE, et al. Antibacterial efficacy of an ultra-short palmitoylated random peptide mixture in mouse models of infection by carbapenem-resistant Klebsiella pneumoniae. Uhlemann AC, editor. Antimicrob Agents Chemother. 2023 Nov 15;67(11):e00574–23.

47. Hayouka Z, Chakraborty S, Liu R, Boersma MD, Weisblum B, Gellman SH. Interplay among Subunit Identity, Subunit Proportion, Chain Length, and Stereochemistry in the Activity Profile of Sequence-Random Peptide Mixtures. J Am Chem Soc. 2013 Aug 14;135(32):11748–51.

48. Hedegaard SF, Derbas MS, Lind TK, Kasimova MR, Christensen MV, Michaelsen MH, et al. Fluorophore labeling of a cell-penetrating peptide significantly alters the mode and degree of biomembrane interaction. Sci Rep. 2018 Apr 20;8(1):6327.

49. Cho J, Lee DG. The antimicrobial peptide arenicin-1 promotes generation of reactive oxygen species and induction of apoptosis. Biochimica et Biophysica Acta (BBA) – General Subjects. 2011 Dec;1810(12):1246–51.

50. Wenzel M, Chiriac AI, Otto A, Zweytick D, May C, Schumacher C, et al. Small cationic antimicrobial peptides delocalize peripheral membrane proteins. Proc Natl Acad Sci USA [Internet]. 2014 Apr 8 [cited 2026 Jan 11];111(14). Available from: https://pnas.org/doi/full/10.1073/pnas.1319900111

51. Maron B, Rolff J, Friedman J, Hayouka Z. Antimicrobial Peptide Combination Can Hinder Resistance Evolution. Sionov E, editor. Microbiol Spectr. 2022 Aug 31;10(4):e00973–22.

52. Yakir I, Cohen E, Schlesinger S, Hayouka Z. Random antimicrobial peptide mixtures as non-antibiotic antimicrobial agents for cultured meat industry. Food Chemistry: Molecular Sciences. 2025 June;10:100240.

53. Gerstein AC, Berman J. Candida albicans Genetic Background Influences Mean and Heterogeneity of Drug Responses and Genome Stability during Evolution in Fluconazole. Mitchell AP, editor. mSphere. 2020 June 24;5(3):e00480–20.

54. Yekani M, Azargun R, Sharifi S, Nabizadeh E, Nahand JS, Ansari NK, et al. Collateral sensitivity: An evolutionary trade-off between antibiotic resistance mechanisms, attractive for dealing with drug-resistance crisis. Health Science Reports. 2023 July;6(7):e1418.

55. Munck C, Gumpert HK, Wallin AIN, Wang HH, Sommer MOA. Prediction of resistance development against drug combinations by collateral responses to component drugs. Sci Transl Med [Internet]. 2014 Nov 12 [cited 2026 Jan 11];6(262). Available from: https://www.science.org/doi/10.1126/scitranslmed.3009940

56. Lara-Aguilar V, Rueda C, García-Barbazán I, Varona S, Monzón S, Jiménez P, et al. Adaptation of the emerging pathogenic yeast *Candida auris* to high caspofungin concentrations correlates with cell wall changes. Virulence. 2021 Dec 31;12(1):1400–17.

57. Mun SH, Joung DK, Kim YS, Kang OH, Kim SB, Seo YS, et al. Synergistic antibacterial effect of curcumin against methicillin-resistant Staphylococcus aureus. Phytomedicine. 2013 June;20(8–9):714–8.

58. Sherry L, Ramage G, Kean R, Borman A, Johnson EM, Richardson MD, et al. Biofilm-Forming Capability of Highly Virulent, Multidrug-Resistant *Candida auris*. Emerg Infect Dis. 2017 Feb;23(2):328–31.

59. Bouza E, Guinea J, Guembe M. The Role of Antifungals against Candida Biofilm in Catheter-Related Candidemia. Antibiotics. 2014 Dec 25;4(1):1–17.

60. Lara HH, Ixtepan-Turrent L, Jose Yacaman M, Lopez-Ribot J. Inhibition of *Candida auris* Biofilm Formation on Medical and Environmental Surfaces by Silver Nanoparticles. ACS Appl Mater Interfaces. 2020 May 13;12(19):21183–91.

61. Horton MV, Nett JE. Candida auris Infection and Biofilm Formation: Going Beyond the Surface. Curr Clin Micro Rpt. 2020 Sept;7(3):51–6.

62. Horton MV, Johnson CJ, Kernien JF, Patel TD, Lam BC, Cheong JZA, et al. Candida auris Forms High-Burden Biofilms in Skin Niche Conditions and on Porcine Skin. Mitchell AP, editor. mSphere. 2020 Feb 26;5(1):e00910–19.

63. Romera D, Aguilera-Correa JJ, Gadea I, Viñuela-Sandoval L, García-Rodríguez J, Esteban J. Candida auris: a comparison between planktonic and biofilm susceptibility to antifungal drugs. Journal of Medical Microbiology. 2019 Sept 1;68(9):1353–8.

64. Kean R, Delaney C, Sherry L, Borman A, Johnson EM, Richardson MD, et al. Transcriptome Assembly and Profiling of Candida auris Reveals Novel Insights into Biofilm-Mediated Resistance. Mitchell AP, editor. mSphere. 2018 Aug 29;3(4):e00334–18.

65. Caraway HE, Lau JZ, Maron B, Oh MW, Belo Y, Brill A, et al. Antimicrobial Random Peptide Mixtures Eradicate Acinetobacter baumannii Biofilms and Inhibit Mouse Models of Infection. Antibiotics. 2022 Mar 19;11(3):413.

66. Zhao M, Lepak AJ, VanScoy B, Bader JC, Marchillo K, Vanhecker J, et al. In Vivo Pharmacokinetics and Pharmacodynamics of APX001 against Candida spp. in a Neutropenic Disseminated Candidiasis Mouse Model. Antimicrob Agents Chemother. 2018 Apr;62(4):e02542–17.

67. Organisation mondiale de la santé, editor. Antimicrobial resistance: global report on surveillance. Genève: World health organization; 2014.

68. Laniado-Laborín R, Cabrales-Vargas MN. Amphotericin B: side effects and toxicity. Revista Iberoamericana de Micología. 2009 Oct;26(4):223–7.

69. Kordalewska M, Lee A, Park S, Berrio I, Chowdhary A, Zhao Y, et al. Understanding Echinocandin Resistance in the Emerging Pathogen Candida auris. Antimicrob Agents Chemother. 2018 June;62(6):e00238–18.

70. Boix-Lemonche G, Lekka M, Skerlavaj B. A Rapid Fluorescence-Based Microplate Assay to Investigate the Interaction of Membrane Active Antimicrobial Peptides with Whole Gram-Positive Bacteria. Antibiotics. 2020 Feb 19;9(2):92.

71. Gbala ID, Macharia RW, Bargul JL, Magoma G. Membrane Permeabilization and Antimicrobial Activity of Recombinant Defensin-d2 and Actifensin against Multidrug-Resistant Pseudomonas aeruginosa and Candida albicans. Molecules. 2022 July 6;27(14):4325.

72. Pavela O, Juhász T, Tóth L, Czajlik A, Batta G, Galgóczy L, et al. Mapping of the Lipid-Binding Regions of the Antifungal Protein NFAP2 by Exploiting Model Membranes. J Chem Inf Model. 2024 Aug 26;64(16):6557–69.

73. Rella A, Farnoud AM, Del Poeta M. Plasma membrane lipids and their role in fungal virulence. Progress in Lipid Research. 2016 Jan;61:63–72.

74. Weghuber J, Aichinger MC, Brameshuber M, Wieser S, Ruprecht V, Plochberger B, et al. Cationic amphipathic peptides accumulate sialylated proteins and lipids in the plasma membrane of eukaryotic host cells. Biochimica et Biophysica Acta (BBA) – Biomembranes. 2011 Oct;1808(10):2581–90.

75. Szomek M, Akkerman V, Lauritsen L, Walther HL, Juhl AD, Thaysen K, et al. Ergosterol promotes aggregation of natamycin in the yeast plasma membrane. Biochimica et Biophysica Acta (BBA) – Biomembranes. 2024 Oct;1866(7):184350.

76. Alavizargar A, Keller F, Wedlich-Söldner R, Heuer A. Effect of Cholesterol Versus Ergosterol on DPPC Bilayer Properties: Insights from Atomistic Simulations. J Phys Chem B. 2021 July 22;125(28):7679–90.

77. Guo L, Li W, Gu Z, Wang L, Guo L, Ma S, et al. Recent Advances and Progress on Melanin: From Source to Application. IJMS. 2023 Feb 22;24(5):4360.

78. Malanovic N, Lohner K. Gram-positive bacterial cell envelopes: The impact on the activity of antimicrobial peptides. Biochimica et Biophysica Acta (BBA) – Biomembranes. 2016 May;1858(5):936–46.

79. Kean R, Ramage G. Combined Antifungal Resistance and Biofilm Tolerance: the Global Threat of Candida auris. Mitchell AP, editor. mSphere. 2019 Aug 28;4(4):e00458–19.

80. Ross ZK, Alsayegh S, Zhao Y, Munro CA, Lorenz A. In vitro evolution of caspofungin resistance in Candidozyma auris via FKS1 hotspot I mutations results in moderate fitness trade-offs but no reduction in virulence [Internet]. 2025 [cited 2026 Jan 13]. Available from: http://biorxiv.org/lookup/doi/10.1101/2024.12.18.629118

81. Carolus H, Pierson S, Muñoz JF, Subotić A, Cruz RB, Cuomo CA, et al. Genome-Wide Analysis of Experimentally Evolved Candida auris Reveals Multiple Novel Mechanisms of Multidrug Resistance. Chowdhary A, editor. mBio. 2021 Apr 27;12(2):e03333–20.

82. Huang X, Chen M, Chen Z, Zhang Y. Fungal β-1,3-glucan synthase: a review of structure, mechanism, and regulation. Munro C, editor. FEMS Yeast Research. 2025 Jan 30;25:foaf071.

83. Dickwella Widanage MC, Gautam I, Sarkar D, Mentink-Vigier F, Vermaas JV, Ding SY, et al. Adaptative survival of Aspergillus fumigatus to echinocandins arises from cell wall remodeling beyond β−1,3-glucan synthesis inhibition. Nat Commun. 2024 July 31;15(1):6382.

84. Wagner AS, Lumsdaine SW, Mangrum MM, Reynolds TB. Caspofungin-induced β(1,3)-glucan exposure in Candida albicans is driven by increased chitin levels. mBio. 2023 Aug 31;14(4):e0007423.

85. Wesseling CMJ, Martin NI. Synergy by Perturbing the Gram-Negative Outer Membrane: Opening the Door for Gram-Positive Specific Antibiotics. ACS Infect Dis. 2022 Sept 9;8(9):1731–57.

86. Spitzer M, Robbins N, Wright GD. Combinatorial strategies for combating invasive fungal infections. Virulence. 2017 Feb 17;8(2):169–85.

87. Sun C, Zhu L, Yang L, Tian Z, Jiao Z, Huang M, et al. Antimicrobial peptide AMP-17 induces protection against systemic candidiasis and interacts synergistically with fluconazole against Candida albicans biofilm. Front Microbiol. 2024 Nov 1;15:1480808.

88. Sun C, Zhao X, Jiao Z, Peng J, Zhou L, Yang L, et al. The Antimicrobial Peptide AMP-17 Derived from Musca domestica Inhibits Biofilm Formation and Eradicates Mature Biofilm in Candida albicans. Antibiotics. 2022 Oct 25;11(11):1474.

89. De La Fuente-Núñez C, Korolik V, Bains M, Nguyen U, Breidenstein EBM, Horsman S, et al. Inhibition of Bacterial Biofilm Formation and Swarming Motility by a Small Synthetic Cationic Peptide. Antimicrob Agents Chemother. 2012 May;56(5):2696–704.

90. Overhage J, Campisano A, Bains M, Torfs ECW, Rehm BHA, Hancock REW. Human Host Defense Peptide LL-37 Prevents Bacterial Biofilm Formation. Infect Immun. 2008 Sept;76(9):4176–82.

91. Zerweck J, Strandberg E, Kukharenko O, Reichert J, Bürck J, Wadhwani P, et al. Molecular mechanism of synergy between the antimicrobial peptides PGLa and magainin 2. Sci Rep. 2017 Oct 13;7(1):13153.

92. Taheri-Araghi S. Synergistic action of antimicrobial peptides and antibiotics: current understanding and future directions. Front Microbiol. 2024 July 31;15:1390765.

93. Cacciapuoti A, Gurnani M, Halpern J, Norris C, Patel R, Loebenberg D. Interaction between Posaconazole and Amphotericin B in Concomitant Treatment against *Candida albicans* In Vivo. Antimicrob Agents Chemother. 2005 Feb;49(2):638–42.

94. Lundvik H, Santini R, Altintop TU, Özenci V, Andersson DI, Fatsis-Kavalopoulos N. Fast detection of synergy and antagonism in antifungal combinations used against Candida albicans clinical isolates. Sci Rep. 2025 Oct 15;15(1):36103.

95. Westerman BA, Broersma Y, Wurdinger T, Noske D, Sminia P, Tannous B. Challenges to determine synergistic drug interactions in mice. Nat Commun. 2025 Sept 30;16(1):8533.

96. Bidaud AL, Schwarz P, Herbreteau G, Dannaoui E. Techniques for the Assessment of In Vitro and In Vivo Antifungal Combinations. JoF. 2021 Feb 4;7(2):113.

97. Zheng S, Tu Y, Li B, Qu G, Li A, Peng X, et al. Antimicrobial peptide biological activity, delivery systems and clinical translation status and challenges. J Transl Med. 2025 Mar 7;23(1):292.

98. Chen SP, Chen EHL, Yang SY, Kuo PS, Jan HM, Yang TC, et al. A Systematic Study of the Stability, Safety, and Efficacy of the de novo Designed Antimicrobial Peptide PepD2 and Its Modified Derivatives Against Acinetobacter baumannii. Front Microbiol. 2021 June 18;12:678330.

